# Structure-based screening of binding affinities via small-angle X-ray scattering

**DOI:** 10.1101/715193

**Authors:** P. Chen, P. Masiewicz, K. Perez, J. Hennig

## Abstract

Protein-protein and protein-ligand interactions can alter the scattering properties of participating molecules, and thus be quantified by solution small-angle X-ray scattering (SAXS). In such cases, scattering reveals structural details of the bound complex, number of species involved, and in principle strength of the interaction. However, determining binding affinities from SAXS-based titrations is not yet an established procedure with well-defined performance expectations. We thus used periplasmic binding proteins and in particular histidine-binding protein as a standard reference, then examined precision and accuracy of affinity prediction at multiple concentrations and exposure times. By analyzing several structural and comparative scattering metrics, we found that the volatility of ratio between titrated scattering curves and a common reference most reliably quantifies ligand-triggered changes. This ratio permits the determination of affinities at low signal-to-noise ratios and without pre-determining the complex scattering, demonstrating that SAXS-based ligand screening is a promising alternative biophysical method for drug discovery pipelines.

**SIGNIFICANCE:** Solution X-ray scattering can be used to screen a set of biomolecular interactions, which yields quantitative information on both structural changes and dissociation constants between binding partners. However, no common benchmarks yet exist for the application of SAXS within drug discovery workflows. Thus, investigations into its performance limitations are currently needed to make SAXS a reliable source for high-throughput screening. This study establishes a generalizable protocol based on protein-ligand interactions, and demonstrates its reproducibility across several beamline setups. In the simplest case, the micromolar binding affinities can be determined directly from measured intensities without knowledge of the molecular structure, with material consumption that is competitive with other biophysical screening techniques.

## INTRODUCTION

Small-angle X-ray scattering (SAXS) is a widely used technique to examine structural features on the micrometer and nanometer scales, offering ready access to the physical behaviours of biomolecules in solution environment.(1–3) Within this context, SAXS reports the globally-averaged distance distribution between scattering electron densities around all atoms. This distribution is obtained via measuring the excess intensity of a sample solution over that of an equivalent buffer solution without sample. The scattered photon intensity *I*(*q*) as a function of the momentum transfer *q*, and its associated scattering angle 2*θ* between the incident and deflected beams, obeys:

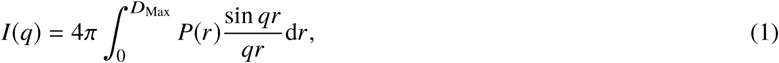

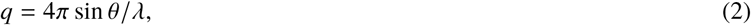

where *P*(*r*) is the desired distance distribution between all atoms in and nearby the molecule with maximum spatial diameter *D*_max_, weighted by the excess electron density. The globally-averaged properties of SAXS implies that a mixture of multiple species with negligible long-range spatial interactions will linearly contribute their respective scattering intensities. Thus, by titrating two species at different input concentrations and measuring *I*(*q*) at each point, their populations at equilibrium can in principle be retrieved along with respective structural information. This is the basis of SAXS-based ligand screening.(4, 5) For illustration, consider a simple two-state interaction between a receptor *R* and its ligand *L* that forms a complex *RL*. The concentrations of these three species at equilibrium are governed by a dissociation constant *K*_D_ that describes the strength of the interaction:

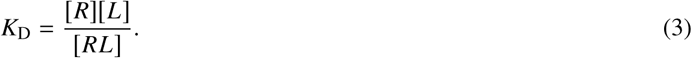

Thus, *K*_D_ can be measured by titrating *R* and *L* at multiple input ligand:receptor ratios. When the size of *L* is small, its scattering is relatively constant over the *q*-ranges covered by SAXS. If we assume that measured *I*(*q*) changes directly correspond to the balance between *R* and *RL* once the constant contribution has been accounted for, then dissociation constants can be directly determined from perturbations of the SAXS signal (Fig. 1).

**Figure 1.**
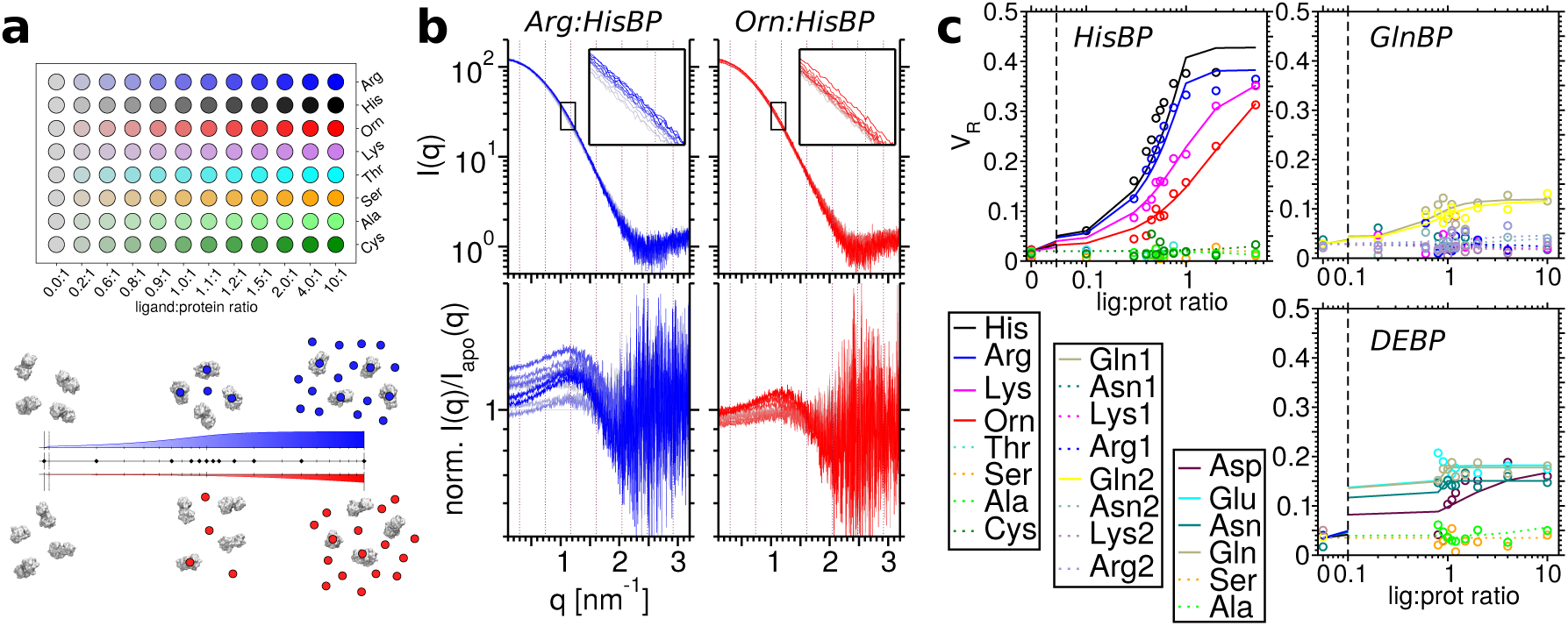
SAXS-based screening using the volatility of ratio *V*_R_, and demonstration on three periplasmic binding proteins: Histidine-binding protein (HisBP), Glutamine-binding protein (GlnBP) and Aspartate/Glutamate-binding protein (DEBP). (a) Many high-throughput synchrotron setups can utilize 96-well plates for small-scale screening, which conveniently fits twelve-point titrations of eight candidates each spanning from the *apo* ligand-less measurement to the saturated ligand-excess measurement. Interactions can be differentiated by tracking the proportion of receptors with a bound ligand. A strong interaction (blue) reaches saturation at approximately 1:1 ligand:protein ratio, whereas a weak interaction (red) may not reach saturation even at 10:1 excess. (b) Titrations of HisBP against Arg (blue) and Orn (red), with minute intensity changes upon ligand saturation. This can be visualized by dividing the intensity *I*(*q*) with the apo-protein *I*(*q*) (pure grey curve), then normalizing the ratio such that its mean is unity. The volatility of ratio *V*_R_ is then computed by averaging data according to the Shannon information content (dotted lines), then summing changes over adjacent channels. Our implementation additionally corrects for unbound ligand contributions by constant subtraction (see SI Methods for details). (c) *V*_R_ values for HisBP (N=96), GlnBP (N=96), and DEBP (N=60) titrations form ligand-dependent binding curves, via which dissociation constants can be computed.

The availability of high-intensity synchrotron sources and automated workflows(6–9) enables precise SAXS measurements within seconds, or far less using time-resolved setups.(10) Thus in the context of screening, a single synchrotron experiment can feasibly screen a small set of candidate ligands and identify members that interact strongly, weakly, or not at all. The precision is such that SAXS can track subtle changes regardless of whether the biomolecules are composed of well-folded domains(11, 12) or disordered chains.(13, 14) This capability has been leveraged to conduct titrations of diverse biomolecular systems: proteins,(15–20) nucleic acids,(21, 22) detergents,(23, 24) and others, in order to expose possible structural mechanisms that underlie their functional behavior.

The majority of above studies concentrate on the structural information obtained, although several so utilize SAXS to additionally compute *K*_D_. Broad uptake of affinity–based applications is hindered by a number of practical difficulties. Firstly, a ligand screen may potentially produce novel complex conformations that are not known prior to the experiment. This complicates the application of population modelling to compute *K*_D_ directly, since the scattering signals of the corresponding purified complex are not available. A reliable, alternative procedure for *K*_D_ predictions must be found using a replacement metric. Secondly, no broadly applicable guidelines have yet been established on how to formulate protocols for SAXS-based ligand screening. While prospective beamline measurements will provide sufficient context to establish sample requirements and measurement protocols, one lacks screening-specific details such as their influence on the observable range and precision of *K*_D_ values. To help tackle this challenge, we conducted reference titrations for a series of bacterial periplasmic binding proteins (PBPs),(25, 26) leveraging their consistent purification and SAXS-detectable compaction upon ligand capture(27) to yield a protocol that can be replicated across multiple beamlines. The availability of SAXS titrations at multiple ligand *K*_D_ and multiple protein concentrations enables us to propose the volatility of ratio *V*_R_ as a viable scattering metric for generalized ligand screening, and begin to define practical sensitivity ranges relevant to SAXS-based ligand screening.

## MATERIALS AND METHODS

### Sample preparation

The *E. coli* HisBP gene (Uniprot: P0AEU0, residues 23-260) was cloned into pET-M11 plasmids containing kanamycin resistance, T7-lac promoter, and an N-terminal His_6_ tag with a TEV cleavage site. The pETMSCIII plasmids for GlnBP (Uniprot: P0AEQ3, residues 23-248) and DEBP (Uniprot: P37902, residues 28-302) were generously provided by Prof. Colin Jackson, containing ampicillin resistance and N-terminal His_6_ tags.

The production and purification of GlnBP and DEBP follow that of previous protocols.(28) We summarize this below with slight modifications for HisBP. After plasmid amplification in *E. coli* DH5*α* cells, protein expression was carried out in *E. coli* BL21(DE3) cells. Production cultures were grown in LB media at 37°C supplemented with either 100 *µ*g/ml ampicillin (GlnBP/DEBP) or 50 *µ*g/ml kanamycin (HisBP). During prospective trials, this was replaced by 15N-labelled minimal media for parallel SAXS, ITC, and NMR experiments. Once OD_600_ exceeded 0.6, cultures were induced with 1 mM isopropyl *β*-D-1-thiogalactopyranoside for 4 hr. Cells were spun-down, resuspended in loading buffer (500 mM NaCl, 20 mM imidazole, and 20 mM NaH_2_PO_4_ [pH 7.4]), lysed by sonication, then filtered before conducting His_6_-tag-based purification by Ni-nitrilotriacetic acid affinity (Ni-NTA) chromatography. After initial loading and washing, on-column refolding was conducted via a decreasing Urea gradient from 8 M to 0 M to remove bound endogenous ligands, then eluted in buffer (500 mM NaCl, 300 mM imidazole, and 20 mM NaH_2_PO_4_ [pH 7.4]). HisBP was further exchanged back into loading buffer, cleaved overnight via addition of TEV protease, then re-loaded onto a second Ni-NTA column to separate the similar-sized N-terminal tag. All PBPs were finally subjected to size-exclusion (SEC) chromatography in the final buffer used for all experimental measurements (100 mM NaCl, 0.5 mM TCEP and 20 mM NaH_2_PO_4_ [pH 7.4]), except for NMR meaurements where samples were diluted due to the addition of 9% D_2_O.

### SAXS measurements

Candidate ligands for each protein were selected on the basis of known affinities in literature and chemical similarity. Titrations were conducted at fixed protein concentrations to exclude signal-to-noise factors. An initial mass concentration of 2.0 or 4.0 mg ml^−1^ was employed in prospective experiments, then scaled to 2.0, 1.0, 0.5, and 0.25 mg ml^−1^ in later concentration variation studies. 12-point titrations were prepared at ligand-protein ratios 0.0, 0.2, 0.6, 0.8, 0.9, 1.0, 1.1, 1.2, 1.5, 20., 4.0, and 10.0 to cover a theoretical *K*_D_ sensitivity within 2 orders of magnitude of the fixed protein concentration employed, assuming ∼5% random error (Fig. S7). A total of eight beamline experiments were conducted, using the automated sample changing environments available on-site, operated in constant-flow mode: three times at ESRF BM29, three at DESY P12, once at Australian Synchrotron SAXS/WAXS, and once at Diamond B21. We note that one prospective session at ESRF and two at DESY have been excluded from this study. Further, experiments at ESRF BM29 and Diamond B21 were carried out on site, while the included DESY P12 experiment was carried out by mail-in. For these measurements, samples were prepared by pipetting ligand:protein mixtures onto 96-well plates prior to transport. As for the Australian Synchrotron measurements, apo-HisBP was lyophilized in its final buffer and reconstituted on-site using pure H_2_O, and then mixed with ligands in 96-well plates. Measurement parameters for each site are catalogued in Table 1. For brevity, we report six representative scattering curves for HisBP, GlnBP, and DEBP in their free and native ligand-bound forms in Table S5 following community guidelines.(29)

**Table 1.** Measurement parameters at the four synchrotron beamlines.

| Parameter | ESRF | DESY | Diamond | AustSynch |
| --- | --- | --- | --- | --- |
| Detector setup | Pilatus-2M | Pilatus-6M | Eiger-4M | Pilatus-1M |
| Wavelength [nm] | 0.9919 | 1.24403 | 0.99987 | 1.07812 |
| $q$-ranged [nm<sup>-1</sup>] | 0.0306–4.9462 | 0.0171–7.2507 | 0.0320–4.3982 | 0.0540–2.8946 |
| Sample-detector distance [m] | 2.849 | 3.0 | 4.014 | 2.68 |
| Per-frame exposure [sec] | 0.5 | 0.1 | 2.0 | 1.0 |
| Frames | 10 | 40 | 20 | 20 |
| Average frame-loss | 0 | 0 | 0 | 3 |
| Sample volume [μl] | 50 | 50 | 50 | 85 |
| Sample temperature [°C] | 20 | 20 | 20 | 20 |

### Automated analysis pipeline

The SAXScreen workflow was utilized for the semi-automated prediction of binding curves from scattering intensities,(20) with modifications for quantitative prediction. Buffer-subtracted intensities provided from respective synchrotron pipelines were used as starting points for analysis. DATGNOM(30) was used to determine a maximum molecular extent *D*_max_ = 7.2 of PBPs, by averaging values found for apo-HisBP at a preliminary ESRF experiment. This was fixed for all computations involving *D*_max_ including *V*_R_. The usable *q*-range was visually determined by conservatively discarding aggregation and noise-dominated regions, and also adjusted to match cross-site data for comparison of similar HisBP concentrations: 0.2–2.0 nm^−1^ below 20 *µ*M, 0.25–2.5 nm^−1^ at 40 *µ*M, and 0.3–3 nm^−1^ above 80 *µ*M. The *q*-range for DEBP and GlnBP was set to 0.1–2.8 nm^−1^. We considered the data cleaning algorithms provided by SAXS Merge to further exclude user bias,(31) but did not use this in the final analysis. A common scattering reference is created per HisBP concentration and per beamline experiment by averaging apo-protein replicates after discarding outliers. This reference is used for both *V*_R_ and χ_lin._ computations. Further explanation of *V*_R_ is available in the Supplementary Material. Other metrics have been computed as previously reported.(20)

To fit binding curves against metrics, we consider as a single set all titrations conducted per protein concentration per beamline experiment. After eliminating measurement artifacts, the set of curves were fitted using a single Powell-minimization algorithm over the following free parameters: the shared apo-value, the holo-values for each ligand titration, and log dissociation affinities log10 *K*_D_ for each ligand titration. Ill-defined regions during fitting are capped by limiting *K*_D_ to within 4 orders of magnitude of the receptor concentration (*c*.*f*. Fig. S7). Two uncertainty estimation methods were implemented to define errorbars in *K*_D_. For metric possessing measurement uncertainty (χ_lin._, *R*_g_ and *V*_c_), 1000 replicates were produced by adding Gaussian-noise with *σ* equal to measurement uncertainty. Error-bars indicate *σ* of log*K*_D_ from the ensemble of replicate predictions. For metrics where uncertainty was not computed (*V*_R_ and PV), the uncertainty was estimated by single-point removal, where *K*_D_ was recomputed after removal of each titration point. The *σ* of the resulting replicates are taken as the *σ* of log*K*_D_. To eliminate cases where the fitting process fails due to ill-constrained parameters, replicates were discarded if the fitted *apo*–*holo* differences were > 3*σ* from the mean value of all replicates.

### ITC

All ITC measurements were conducted on a Malvern MicroCal PEAQ ITC at 20°C with stirring at 200 rpm, using the same buffer as in SAXS measurements. The cell was initially loaded with 200 *µ*l of 20 *µ*M protein with an initial ligand concentration of 200 *µ*M in the syringe to quantify HisBP:His binding. This was further adjusted in subsequent runs to resolve weaker interactions as necessary. ITC titration curves for all experiments are shown in Figure S14 along with the concentrations used.

### NMR

All NMR experiments have been carried out at 298 K on an 800 MHz Bruker Avance III NMR spectrometer, equipped with a cryogenic probehead. Titrations of HisBP against His and Arg were measured via recording ^1^*H*-^15^*N* HSQC spectra at discrete ligand:protein-ratios of 0.0, 0.1, 0.2, 0.3, 0.4, 0.6, 0.8, 0.9, 1.0, 1.1, 1.2, and 1.5 for His; and 0.0, 0.1, 0.2, 0.3,0.4, 0.6, 0.8, 0.9, 1.0, 1.1, 1.2, 1.5, 1.8 and 2.1 for Arg, respectively. Data was processed with NMRpipe,(32) analyzed using NMRFAM-Sparky,(33) and visualized with python modules nmrglue(34) and matplotlib.(35)

Backbone assignments of apo- and His-bound HisBP have been derived from BMRB entries 19242 and 19245.(36) The assignment of Arg-bound HisBP is produced by extending the apo-HisBP assignment along titrations, leaving sites in slow exchange unassigned. For affinity computations based on chemical shift perturbations, ^1^*H* and ^15^*N* shifts have been combined to a composite shift 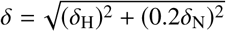. The derivation of an overall dissociation constant from chemical shift perturbations follows Williamson (2013)(37), where the shift in peak position is used to derive free and bound populations without considering peak intensities. The effect of site-specific exchange rates upon peak positions is taken into account.(38) We discarded residues with either insignificant shifts at saturation (*δ* < 0.05), or which were in slow exchange such that peak positions alone were no longer informative. This resulted in 110 sites included in the fitting process. The binding interaction is modeled with site-specific bound-state CSP, site-specific bound-state lifetimes *τ*, and a single overall *K*_D_. This translates to 221 degrees of freedom, fitted to 1468 target CSP values.

### Software and data availability

The SAXScreen software has been previously published and made available via https://www.github.com/zharmad/SAXScreen. The set of sample and buffer 1D-scattering intensities for each beamline will be made available via https://zenodo.org, accessible via the DOI:10.5281/zenodo.3355877. Representative scattering datasets have been deposited at SASBDB(39) encompassing HisBP, DEBP, and GlnBP in their unbound states and bound to eponymous cognate ligands (SASBDB identifiers: SASDFD8, SASDFE8, SASDFF8, SASDFG8, SASDFH8 and SASDFJ8).

## RESULTS

### Evaluation of multiple PBPs at ESRF Grenoble

Within the *in vivo* context, PBPs serve as nutrient sensing and transport proteins, and are physically composed of two hinged globular domains each providing one half of a ligand binding site within their central cleft. We surveyed a set of amino acid specific PBPs with preliminary SAXS measurements conducted at ESRF BM29 and DESY P12 to identify a member that is best suited for evaluating the performance of SAXS-based ligand screening: Histidine-binding protein (HisBP), glutamine-binding protein (GlnBP), and aspartate/glutamate-binding protein (DEBP). In the absence of ligand, PBPs exist either as purely open conformations as is the case for DEBP, or already in a pre-existing open-closed conformational equilibrium in the cases of HisBP and GlnBP.(41) Ligand binding universally increases the population of closed/bound conformations, where the physical compaction is measurable as a small but significant change in the SAXS radius of gyration *R*_g_ : 1.34±0.05 Å for HisBP:His binding, 0.66±0.29Å for GlnBP:Gln binding, and 3.23±0.23Å for DEBP:Glu binding. The net scattering changes can also be summarized via other parameters, such as Porod volume PV, linearity of fit χ_lin._,(20) volume of correlation *V*_c_,(42) and volatility of ratio *V*_R_.(43) *V*_R_ in particular compares the rate of intensity changes between two scattering curves – its nomenclature arises from economics where *V*_R_ is used to compare the “riskiness” between two stocks, *i*.*e*. which is more prone to rapid fluctuations in price. The adoption of *V*_R_ in SAXS serves as a generalized measure of scattering difference that is unbiased over *q* and also insensitive to protein size. To enable comparisons between titrations within a single SAXS experiment, we average the replicate apo-protein scattering curves per concentration and compute *V*_R_ against these averages, and further take into account unbound ligand scattering by adding constant corrections to minimize *V*_R_.

The scattering perturbations of a series of ligands against HisBP, GlnBP, and DEBP at a preliminary ESRF session is summarized in Fig. 1c. The high protein concentrations in this initial test (2 ∼ 4 mg ml^−1^) give rise to excellent signal-to-noise ratios. This was necessary to detect the relatively minute GlnBP perturbations, and in addition limits the deterioration of accuracy due to lost data points. Despite the loss of 0.2:1 and 0.6:1 scattering curves in DEBP titrations, binding could still be quantified. Of the tested candidates, the significant scattering perturbations and availability of both nanomolar and micromolar interactions suggest that HisBP is the best candidate to use for the further exploration of screening protocols.

Aside from the binding curves exhibited in *V*_R_, we also observe analogous binding interactions exhibited by the majority of structural parameters (Fig S1). However, these other parameters appear to exhibit significantly lower precision due to measurement and preparation limitations. While χ_lin_ also corrects for small concentration variations, and appears to be the next-best performing, applying linear fitting to raw intensities appears to over-emphasize the contributions from low angles where signal-to-noise is at its optimum. This renders χ_lin_ vulnerable to residual aggregation artifacts, which also affects the net molecular compactness as measured by *R*_g_. These limitations to sample purity are particularly problematic for GlnBP, where the scattering intensity itself experiences a maximum of only 10% change upon ligand capture (Fig. S2). The overall size as reported by Porod volume is limited by our ability to precisely prepare molecular concentrations, estimated to similarly possess ∼10% error. Only DEBP exhibits an apparent size change greater than this margin. We also note that *V*_c_ appears to pick up universal binding at high ligand saturation inconsistent with all other metrics. This is likely to be a false detection of gradual buffer differences stemming from unbound ligand and counter-ions. Regardless of the metric used, significant artifacts such as from sample loss cannot be corrected *post-hoc* and must be manually removed.

While the scattering perturbations closely resemble binding curves, the derived *K*_D_ values are not always in agreement with nominally equivalent estimates derived from ITC (Table 2). For instance, almost all SAXS titrations estimate the nanomolar HisBP:His and GlnBP:Gln interactions instead to be 10-100 fold weaker. Curiously, SAXS suggests that DEBP binds not only its known ligands Glu and Asp, but also their polar equivalents Gln and Asn. Here, it is important to note that the two sources need not be consistent because SAXS measures structural change while ITC measures binding thermodynamics. SAXS-based *K*_D_ reports the structural equilibrium between bound-closed/free-closed versus both bound-open/free-open conformations. If ligand binding alone is not sufficient to fully stabilize a closed conformation, then this may differ from the ITC measurement, which reports the net heat produced arising from both ligand-binding and conformational change steps. Another explanation for the discrepancy is that the sample requirements in our SAXS protocols are unable to effectively discriminate nanomolar affinities for PBP-sized biomolecules (∼30 kDa). This is explored further below.

**Table 2.** Summary of fitted dissociation constants *K*_D_ derived from *V*_R_ titration curves, with comparisons against ITC and NMR measurements. Titrations that produce no significant changes in scattering curves have been denoted as *no change*.

| Protein | source | exposure | conc.<br>($\mu\text{M}$) | Ligand identity and dissociation constant $K_D$ | | | |
| --- | --- | --- | --- | --- | --- | --- | --- |
|  |  |  |  | His | Arg | Lys | Orn |
| HisBP | $V_R$ (DESY) | 4 s | 80 | $6.4 \pm 1.1 \mu\text{M}$ | $5.9 \pm 0.6 \mu\text{M}$ | $58 \pm 7.5 \mu\text{M}$ | $100 \pm 26 \mu\text{M}$ |
| | $V_R$ (DESY) | 4 s | 20 | $1 \pm 0.19 \mu\text{M}$ | $33 \pm 4.5 \mu\text{M}$ | $61 \pm 6.4 \mu\text{M}$ | $130 \pm 170 \mu\text{M}$ |
| | $V_R$ (ESRF) | 5 s | 160 | $370 \pm 470 \text{ nM}$ | $710 \pm 1000 \text{ nM}$ | $49 \pm 4.1 \mu\text{M}$ | $250 \pm 24 \mu\text{M}$ |
| | $V_R$ (ESRF) | 5 s | 38 | $240 \pm 140 \text{ nM}$ | $2 \pm 1.9 \mu\text{M}$ | n.d. | $5.4 \pm 28 \text{ mM}$ |
| | $V_R$ (AustSynch) | 20 s | 76 | $3.6 \pm 0.74 \mu\text{M}$ | $5.5 \pm 1.4 \mu\text{M}$ | $95 \pm 11 \mu\text{M}$ | $220 \pm 34 \mu\text{M}$ |
| | $V_R$ (AustSynch) | 20 s | 19 | $62 \pm 28 \text{ nM}$ | $2.8 \pm 0.31 \mu\text{M}$ | $110 \pm 10 \mu\text{M}$ | $130 \pm 24 \mu\text{M}$ |
| | $V_R$ (Diamond) | 40 s | 80 | $1.3 \pm 0.5 \mu\text{M}$ | $7.3 \pm 1.9 \mu\text{M}$ | $89 \pm 9.6 \mu\text{M}$ | $640 \pm 160 \mu\text{M}$ |
| | $V_R$ (Diamond) | 40 s | 20 | $\ll 40 \text{ nM}$ | $920 \pm 110 \text{ nM}$ | $47 \pm 3.6 \mu\text{M}$ | $1.4 \pm 1.1 \text{ mM}$ |
| | ITC(40) | – | – | $64 \pm 10 \text{ nM}$ | $3 \pm 1 \mu\text{M}$ | $19.5 \pm 6 \mu\text{M}$ | $72 \pm 4 \mu\text{M}$ |
| | ITC | – | – | $48.0 \pm 5.2 \text{ nM}$ | $2.06 \pm 0.22 \mu\text{M}$ | $62 \pm 11 \mu\text{M}$ | $220 \pm 37 \mu\text{M}$ |
|  | NMR | – | 200 | – | 560 nM |  |  |
| DEBP |  |  |  | Glu | Asp | Gln | Asn |
| | ITC(28) | – | – | $334 \pm 120 \text{ nM}$ | $1.73 \pm 0.49 \mu\text{M}$ | – | – |
| | ITC, this work | – | – | $141 \pm 66 \text{ nM}$ | $1.15 \pm 0.18 \mu\text{M}$ | – | – |
| | $V_R$ (ESRF) | 5 s | 80 | $150 \pm 150 \text{ nM}$ | $63 \pm 34 \mu\text{M}$ | $\ll 80 \text{ nM}$ | $\ll 80 \text{ nM}$ |
| GlnBP |  |  |  | Gln | Arg |  |  |
| | ITC(28) | – | – | $106 \pm 65 \text{ nM}$ | $93.8 \pm 7.4 \mu\text{M}$ | | |
| | ITC | – | – | $59 \pm 37 \text{ nM}$ | $78.9 \pm 8.6 \mu\text{M}$ | | |
| | $V_R$ (ESRF) | 5 s | 80 | $8.2 \pm 7.5 \mu\text{M}$ | *no change* | | |
| | $V_R$ (ESRF) | 5 s | 80 | $18 \pm 10 \mu\text{M}$ | *no change* | | |

### HisBP replication studies

We investigated the sensitivity limits of SAXS-based screening by conducting HisBP titrations at multiple protein concentrations, selected to explore the balance between sample requirements and discriminative power between nanomolar affinities. Resulting *V*_R_ measurements at four beamlines are reported in Fig. 2 with respective *K*_D_ predictions visualized in Fig. 3. A summary of *K*_D_ values is included in Table 2 with full values in Table S1. For completeness, we include results for χ_lin_, *R*_g_, and *V*_c_ in Figures S3–S5 and corresponding *K*_D_ values in Tables S2–S4.

**Figure 2.**
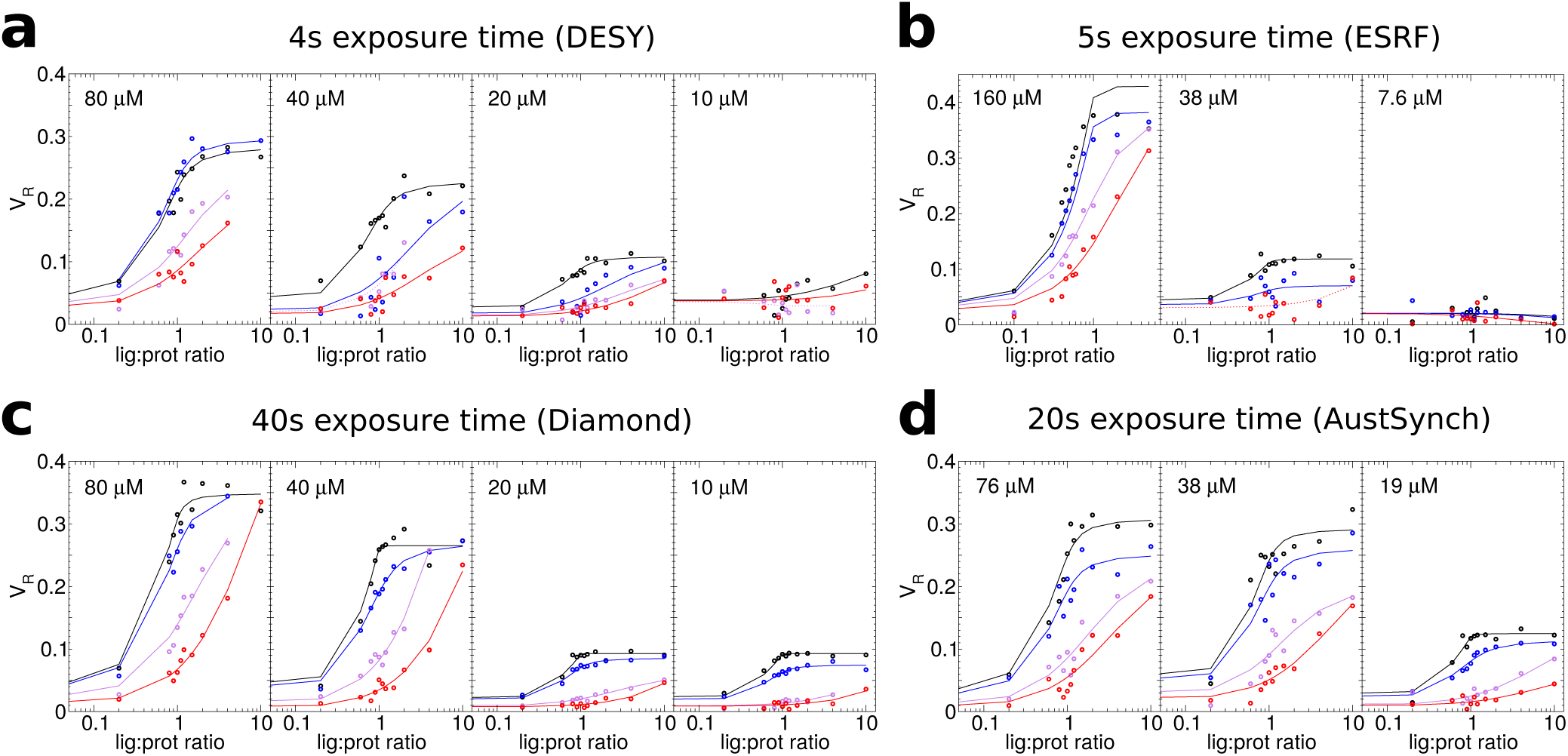
Replicate *V*_R_ titration curves of HisBP against four ligands His (black), Arg (blue), Lys (violet), and Orn (red), collated at DESY (a, N=172), ESRF (b, N=144), Diamond (c, N=192), and Australian Synchrotron (d, N=144). The HisBP concentrations used within each experiment are labelled on respective plots. Note that the maximum *q*-range measured in (d) was 2.8 nm^−1^, which affected the *V*_R_.

**Figure 3.**
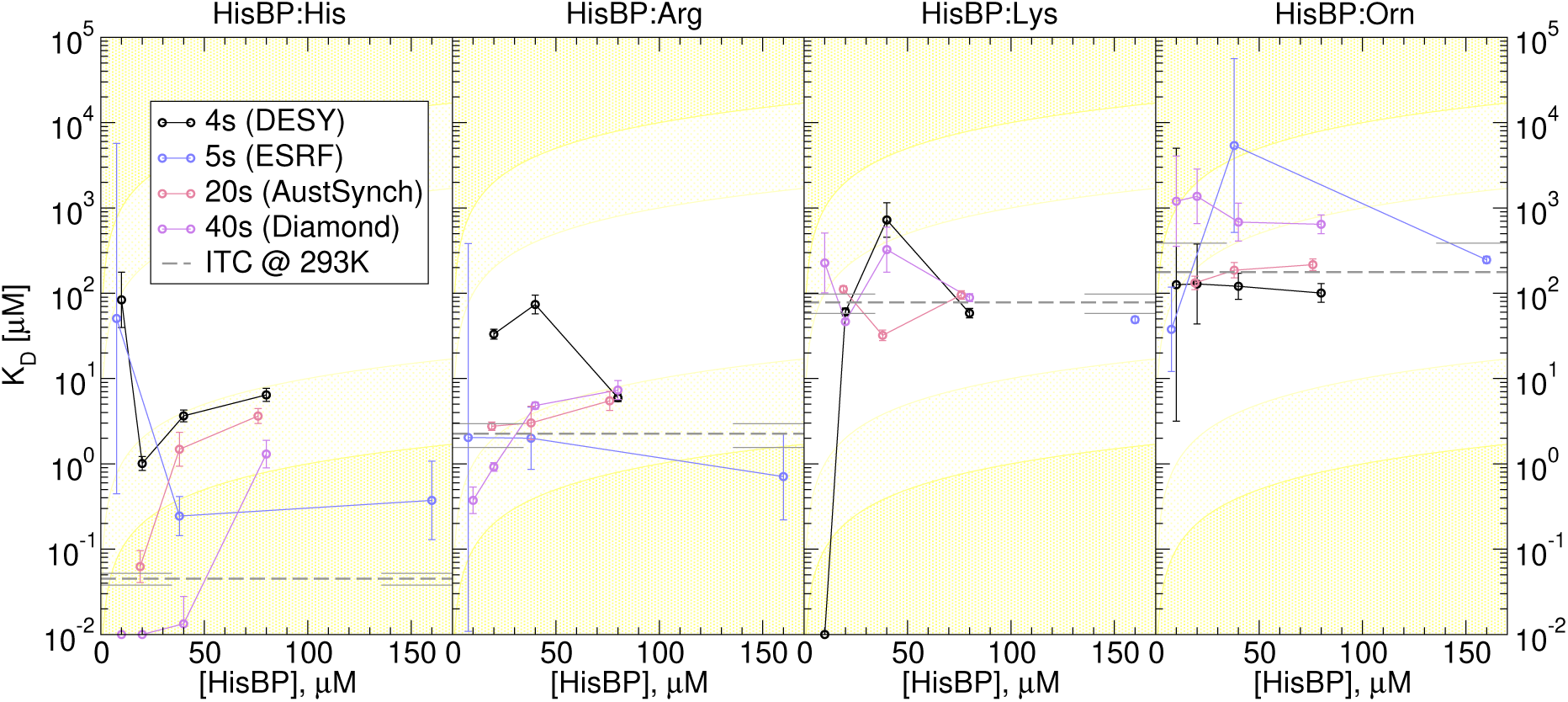
HisBP screens replicated at DESY P12 (4 s exposure, black), ESRF BM29 (5 s exposure. blue), Diamond B21 (40 s exposure, violet) and Australian Synchrotron SAXS/WAXS (20 s exposure. red), using *V*_R_ to derive relative populations. The affinity *K*_D_ of HisBP versus four ligands is evaluated at different protein concentrations and exposure times to test prediction capability versus ITC values (dotted grey with error bars). Regions where *K*_D_ predictions lie outside 1 and 2 orders of magnitude of input protein concentrations are shaded in light and dark yellow, representing regions of decreased confidence.

We first find that the total *V*_R_ change upon ligand saturation decreases with decreasing HisBP concentration, due to the concomitant decrease in usable *q*-range. In the case of HisBP, dropping concentrations from 40 *µ*M to 20 *µ*M lead to loss of detectable intensity changes above *q* > 2 nm^−1^ (Fig. S6). The remaining scattering range is nevertheless sufficient to quantify binding at most concentrations. The minimum viable concentration is imposed by total sample exposure time: the short exposure times of 4–5 s trialled at DESY and ESRF provided insufficient signal-to-noise ratios to quantify scattering changes using 10 *µ*M protein, corresponding to 0.26 mg ml^−1^. In contrast, long 40 s exposures trialled at Diamond enable screening at this concentration, which is near the sample limitations recommended for synchrotron scattering. These findings suggest that trade-off between performance and sensitivity occurs below 20 *µ*M for HisBP. Although exact numbers are also influenced by additional factors such as source intensity and sample stability, we expect comparable limitations for similarly-sized systems exhibiting total *V*_R_ change of 0.1.

Repeated HisBP titrations also show that SAXS-based titrations conform to the *K*_D_ range limits imposed by two-state binding (Fig. S7). In particular, we observe a strong dependence upon protein concentration of the fitted *K*_D_ values for HisBP:His, and to a lesser extent HisBP:Arg (Fig. 3). This systematic error excludes the alternative hypothesis that histidine is unable to fully stabilize a closed conformation, which is also corroborated by observations in NMR chemical shifts of slow kinetics and saturation near 1:1-ratios (Fig. S8). Although we initially estimated that *K*_D_ cannot be distinguished below 2 orders of magnitude of protein concentrations, the practical limit appears to be somewhat closer to 1. Regardless of the quantitative limitations, *V*_R_ retains qualitative ranking between the four ligands for the majority of experiments. Longer exposure protocols appear to provide both more consistent ranking and less severe systematic errors. Further analysis of scattering parameters also suggest structural differences between the His-bound and Arg-bound HisBPs. Both *V*_R_ and χ_lin_ indicate larger perturbations elicited by His binding, in contrast to their comparable total change in *R*_g_. This suggests that HisBP reorients its domains laterally to accommodate the larger Arg sidechain. We further confirmed the results in NMR titrations of the two HisBP ligands: while the active site is shared, the extent and direction of shifts are altered at numerous locations (Fig. S8-S10).

## DISCUSSION

The practical niche for SAXS-based ligand screening appears to be a complementary filter that grants structural information on the bound configurations, providing information on the equilibrium between structurally distinguishable states. This provides a unique advantage in determining whether candidate ligands elicit or inhibit structural changes required for native function, serving as an *in vitro* discriminator between agonists and antagonists in the context of drug discovery. When permitted by sample stability and available beam time, it is possible to quantitatively derive *K*_D_ at the lower sample limits of synchrotron scattering, giving SAXS a competitive advantage on this aspect versus alternative structural biology methods. On the other hand, the ability to differentiate between strong interactions is limited by minimum receptor concentrations required to obtain scattering data and detect potentially minute variations. Outside of large (> 100 kDa) complexes exhibiting significant global rearrangements, we do not expect SAXS to provide nanomolar resolution. While the tested scattering parameters all provide information on the mixture of states during titration, *V*_R_ appears to be the most reliable method to extract the underlying populations and thus a means of quantifying *K*_D_. Notably, no assumption is made of the substrate material. It is possible that *V*_R_ can quantify structural alterations in other existing applications of SAXS, for instance cellular ultrastructure,(44) whole-cell morphology,(45) and beyond.

The HisBP titrations conducted here are sufficient to provide initial guidelines on viable SAXS-based ligand screening protocols. A 20 *µ*M limit for HisBP translates to a minimum sample requirement of 0.31 mg per titration, competitive with ∼0.24 mg used per ITC run and 2.4 mg used for NMR titration. This is again likely to hold for similarly-sized systems exhibiting comparable structural perturbations. The minimum consumption is ultimately determined by the replicate measurements necessary to reliably determine apo, 1:1, and ligand-excess scattering. This is in turn determined by *accuracy* of *V*_R_ measurements versus its expected change due to structural perturbations. We note that *V*_R_ of independent apo measurements constitutes a measure of accuracy, as its ideal value is zero. In contrast, the expected *V*_R_ changes due to binding is system-dependent. Further work will be needed to derive an algorithm that predicts the expected *V*_R_ changes based on the receptor structure or scattering pattern, and thus determine screening viability. This will also help determine whether *V*_R_ is a universally applicable metric for affinity computation, both in terms of the type of structural changes and in terms of reaction complexity. It is possible that other metrics may be superior in cases where intensity changes are restricted to particular *q*-regions, or where more than two states are involved. Here, we expect that direct population modelling of intensities will persist as the theoretically optimal choice.

The throughput performance of each beamline used here varies between 3.7 hours and 6 hours per 96-well plate depending on overall exposure time, sample handling, and cleaning of measurement capillaries. The last two are major limiting factor to further improvement for solution setups, where the next breakthroughs in both speed and precision are likely to take place on existing robotic and promising microfluidic-chip(46–48) platforms. Although not employed here, we note that the SIBYLS beamline claims highest throughput so far at ∼15 minutes per plate with unit second exposures. Adjusting for 20 s exposure times required to improve precision, this decays to ∼45 minutes per plate, sufficient to screen 10^2^ ligands in a 24-hour session. For reference, our ITC and NMR protocols require ∼48 hours to accumulate 96 measurements. This is significantly slower, but is less constrained by available access.

## CONCLUSIONS

In summary, this work presents a detailed analysis of the accuracy and precision of SAXS-based *K*_D_ determination, using sample setups that are competitive with secondary screening approaches used to complement high-throughput screening. In comparison to the throughput of pharmacological screening assays (up to 10^6^ compounds,) the throughput here is sufficient to validate a selection of initial hits and inform on the structural implications of ligand binding. This translates to discrimination between likely agonists and antagonists, potential oligomerization, and other observations that may not be available from standard pharmacological. In this way, SAXS-based ligand screening can benefit drug discovery pipelines as an independent source of structural information for solution biological processes, without requiring isotope-labelling for NMR or reliable crystallization protocols. Further work will be needed to confirm that *V*_R_ can be used to directly retrieve *K*_D_ regardless of the biological system being used, which will also contribute towards a useful library of reference data to guide future screening efforts. Along with potential feasibility studies using lab-based X-ray sources, we may yet see a mass uptake of this particular structure-based screening tool.

## AUTHOR CONTRIBUTIONS

PC and JH conceived and designed the research. PC conducted NMR and the majority of SAXS measurements, analyzed data, and updated SAXScreen software. PM conducted half of sample preparations and trained PC to conduct the other half. KP conducted the majority of ITC measurements and trained PC for the remaining measurements, as well as analysis. JH conducted two of the eight SAXS measurements. PC and JH wrote the article, which was reviewed by all authors.

## ACKNOWLEDGEMENTS

We gratefully acknowledge the provision of beam time from the following synchrotron facilities and support from their respective staff: Petra Pernot at ESRF BM29; Karen Manalastas, Tobias Graewert, Melissa Anne Graewert, and Nelly Hajizadeh at DESY P12; Nathan Cowieson at Diamond B21; and Tim Ryan at the Australian Synchrotron SAXS/WAXS. We also thank Prof. Dmitri Svergun for valuable discussions during initial screening trials at DESY P12, and Prof. Colin Jackson for the provision of GlnBP and DEBP plasmids. P.C. was supported by the EIPOD postdoctoral programme co-funded by the European Molecular Biology Laboratory (EMBL) and the Marie Curie Actions Cofund grant MSCA-COFUND-FP #664726. J.H. gratefully acknowledges the EMBL and the German Research Council (Deutsche Forschungsgemeinschaft, DFG) for support via an Emmy-Noether Fellowship.

## REFERENCES

1. Svergun, D. I., M. H. J. Koch, P. A. Timmins, and R. P. May, 2013. Small Angle X-Ray and Neutron Scattering from Solutions of Biological Macromolecules. Oxford University Press.

2. Putnam, C. D., M. Hammel, G. L. Hura, and J. A. Tainer, 2007. X-Ray Solution Scattering (SAXS) Combined with Crystallography and Computation: Defining Accurate Macromolecular Structures, Conformations and Assemblies in Solution. Q. Rev. Biophys. 40:191–285.

3. Skou, S., R. E. Gillilan, and N. Ando, 2014. Synchrotron-Based Small-Angle X-Ray Scattering of Proteins in Solution. Nat. Protocols 9:1727–1739.

4. Tuukkanen, A. T., and D. I. Svergun, 2014. Weak Protein–Ligand Interactions Studied by Small-Angle X-Ray Scattering. FEBS J 281:1974–1987.

5. Chen, P.-c., and J. Hennig, 2018. The Role of Small-Angle Scattering in Structure-Based Screening Applications. Biophys.Rev. 10:1295–1310.

6. Classen, S., G. L. Hura, J. M. Holton, R. P. Rambo, I. Rodic, P. J. McGuire, K. Dyer, M. Hammel, G. Meigs, K. A. Frankel, and J. A. Tainer, 2013. Implementation and Performance of SIBYLS: A Dual Endstation Small-Angle X-Ray Scattering and Macromolecular Crystallography Beamline at the Advanced Light Source. J. Appl. Crystallogr. 46:1–13.

7. Kirby, N. M., S. T. Mudie, A. M. Hawley, D. J. Cookson, H. D. T. Mertens, N. Cowieson, and V. Samardzic-Boban, 2013. A Low-Background-Intensity Focusing Small-Angle X-Ray Scattering Undulator Beamline. J. Appl. Crystallogr. 46:1670–1680.

8. Acerbo, A. S., M. J. Cook, and R. E. Gillilan, 2015. Upgrade of MacCHESS Facility for X-Ray Scattering of Biological Macromolecules in Solution. J. Synchrotron Radiat. 22:180–186.

9. Blanchet, C. E., A. Spilotros, F. Schwemmer, M. A. Graewert, A. Kikhney, C. M. Jeffries, D. Franke, D. Mark, R. Zengerle, F. Cipriani, S. Fiedler, M. Roessle, and D. I. Svergun, 2015. Versatile Sample Environments and Automation for Biological Solution X-Ray Scattering Experiments at the P12 Beamline (PETRA III, DESY). J. Appl. Crystallogr. 48:431–443.

10. Cammarata, M., M. Levantino, F. Schotte, P. A. Anfinrud, F. Ewald, J. Choi, A. Cupane, M. Wulff, and H. Ihee, 2008. Tracking the Structural Dynamics of Proteins in Solution Using Time-Resolved Wide-Angle X-Ray Scattering. Nat. Methods 5:881–886.

11. Rambo, R. P., and J. A. Tainer, 2013. Super-Resolution in Solution X-Ray Scattering and Its Applications to Structural Systems Biology. Ann. Rev. Biophys. 42:415–441.

12. Vestergaard, B., and Z. Sayers, 2014. Investigating Increasingly Complex Macromolecular Systems with Small-Angle X-Ray Scattering. IUCrJ 1:523–529.

13. Kikhney, A. G., and D. I. Svergun, 2015. A Practical Guide to Small Angle X-Ray Scattering (SAXS) of Flexible and Intrinsically Disordered Proteins. FEBS Lett. 589:2570–2577.

14. Cordeiro, T. N., F. Herranz-Trillo, A. Urbanek, A. Estaña, J. Cortés, N. Sibille, and P. Bernadó, 2017. Structural Characterization of Highly Flexible Proteins by Small-Angle Scattering. In Biological Small Angle Scattering: Techniques, Strategies and Tips, Springer, Singapore, Advances in Experimental Medicine and Biology, 107–129.

15. Ando, N., Y. Kung, M. Can, G. Bender, S. W. Ragsdale, and C. L. Drennan, 2012. Transient B12-Dependent Methyltransferase Complexes Revealed by Small-Angle X-Ray Scattering. J. Am. Chem. Soc. 134:17945–17954.

16. Ando, N., H. Li, E. J. Brignole, S. Thompson, M. I. McLaughlin, J. E. Page, F. J. Asturias, J. Stubbe, and C. L. Drennan, 2016. Allosteric Inhibition of Human Ribonucleotide Reductase by dATP Entails the Stabilization of a Hexamer. Biochemistry 55:373–381.

17. Cordeiro, T. N., P.-c. Chen, A. De Biasio, N. Sibille, F. J. Blanco, J. S. Hub, R. Crehuet, and P. Bernadó, 2017. Disentangling Polydispersity in the PCNA-p15PAF Complex, a Disordered, Transient and Multivalent Macromolecular Assembly. Nucl. Acids Res. 45:1501–1515.

18. Fukuda, M., A. Watanabe, A. Hayasaka, M. Muraoka, Y. Hori, T. Yamazaki, Y. Imaeda, and A. Koga, 2017. Small-Scale Screening Method for Low-Viscosity Antibody Solutions Using Small-Angle X-Ray Scattering. Eur. J. Pharmaceut. Biopharmaceut. 112:132–137.

19. Tian, X., A. E. Langkilde, M. Thorolfsson, H. B. Rasmussen, and B. Vestergaard, 2014. Small-Angle X-Ray Scattering Screening Complements Conventional Biophysical Analysis: Comparative Structural and Biophysical Analysis of Monoclonal Antibodies IgG1, IgG2, and IgG4. J. Pharmaceut. Sci. 103:1701–1710.

20. Chen, P.-c., P. Masiewicz, V. Rybin, D. Svergun, and J. Hennig, 2018. A General Small-Angle X-Ray Scattering-Based Screening Protocol Validated for Protein–RNA Interactions. ACS Comb. Sci. 20:197–202.

21. Meisburger, S. P., J. L. Sutton, H. Chen, S. A. Pabit, S. Kirmizialtin, R. Elber, and L. Pollack, 2013. Polyelectrolyte Properties of Single Stranded DNA Measured Using SAXS and Single-molecule FRET: Beyond the Wormlike Chain Model. Biopolymers 99:1032–1045.

22. Chen, Y., and L. Pollack, 2016. SAXS Studies of RNA: Structures, Dynamics, and Interactions with Partners. WIREs RNA 7:512–526.

23. Lipfert, J., L. Columbus, V. B. Chu, S. A. Lesley, and S. Doniach, 2007. Size and Shape of Detergent Micelles Determined by Small-Angle X-Ray Scattering. J. Phys. Chem. B 111:12427–12438.

24. Oliver, R. C., J. Lipfert, D. A. Fox, R. H. Lo, S. Doniach, and L. Columbus, 2013. Dependence of Micelle Size and Shape on Detergent Alkyl Chain Length and Head Group. PLoS ONE 8:e62488.

25. Tam, R., and M. H. Saier, 1993. Structural, Functional, and Evolutionary Relationships among Extracellular Solute-Binding Receptors of Bacteria. Microbiol. Rev. 57:320–346.

26. Quiocho, F. A., and P. S. Ledvina, 1996. Atomic Structure and Specificity of Bacterial Periplasmic Receptors for Active Transport and Chemotaxis: Variation of Common Themes. Mol. Microbiol. 20:17–25.

27. Olah, G. A., S. Trakhanov, J. Trewhella, and F. A. Quiocho, 1993. Leucine/Isoleucine/Valine-Binding Protein Contracts upon Binding of Ligand. J. Biol. Chem. 268:16241–16247.

28. Clifton, B. E., and C. J. Jackson, 2016. Ancestral Protein Reconstruction Yields Insights into Adaptive Evolution of Binding Specificity in Solute-Binding Proteins. Cell Chem. Biol. 23:236–245.

29. Trewhella, J., A. P. Duff, D. Durand, F. Gabel, J. M. Guss, W. A. Hendrickson, G. L. Hura, D. A. Jacques, N. M. Kirby, H. Kwan, J. Pérez, L. Pollack, T. M. Ryan, A. Sali, D. Schneidman-Duhovny, T. Schwede, D. I. Svergun, M. Sugiyama, J. A. Tainer, P. Vachette, J. Westbrook, and A. E. Whitten, 2017. 2017 Publication Guidelines for Structural Modelling of Small-Angle Scattering Data from Biomolecules in Solution: An Update. Acta Crystallogr. D 73:710–728.

30. Franke, D., M. V. Petoukhov, P. V. Konarev, A. Panjkovich, A. Tuukkanen, H. D. T. Mertens, A. G. Kikhney, N. R. Hajizadeh, J. M. Franklin, C. M. Jeffries, and D. I. Svergun, 2017. ATSAS 2.8: A Comprehensive Data Analysis Suite for Small-Angle Scattering from Macromolecular Solutions. J. Appl. Crystallogr. 50:1212–1225.

31. Spill, Y. G., S. J. Kim, D. Schneidman-Duhovny, D. Russel, B. Webb, A. Sali, and M. Nilges, 2014. SAXS Merge: An Automated Statistical Method to Merge SAXS Profiles Using Gaussian Processes. J. Synchrotron Radiat. 21:203–208.

32. Delaglio, F., S. Grzesiek, G. W. Vuister, G. Zhu, J. Pfeifer, and A. Bax, 1995. NMRPipe: A Multidimensional Spectral Processing System Based on UNIX Pipes. J. Biomol. NMR 6:277–293.

33. Lee, W., M. Tonelli, and J. L. Markley, 2015. NMRFAM-SPARKY: Enhanced Software for Biomolecular NMR Spectroscopy. Bioinformatics 31:1325–1327.

34. Helmus, J. J., and C. P. Jaroniec, 2013. Nmrglue: An Open Source Python Package for the Analysis of Multidimensional NMR Data. J. Biomol. NMR 55:355–367.

35. Hunter, J. D., 2007. Matplotlib: A 2D Graphics Environment. Comput. Sci. Eng. 9:90–95.

36. Chu, B. C. H., T. DeWolf, and H. J. Vogel, 2013. Role of the Two Structural Domains from the Periplasmic Escherichia Coli Histidine-Binding Protein HisJ. J. Biol. Chem. 288:31409–31422.

37. Williamson, M. P., 2013. Using Chemical Shift Perturbation to Characterise Ligand Binding. Prog. Nucl. Magn. Reson.Spectrosc. 73:1–16.

38. London, R. E., 1993. Chemical-Shift and Linewidth Characteristics of Reversibly Bound Ligands. J. Magn. Reson. 104:190–196. London. JMagnReson.1993.

39. Valentini, E., A. G. Kikhney, G. Previtali, C. M. Jeffries, and D. I. Svergun, 2015. SASBDB, a Repository for Biological Small-Angle Scattering Data. Nucl. Acids Res. 43:D357–D363.

40. Paul, S., S. Banerjee, and H. J. Vogel, 2016. Ligand Binding Specificity of the Escherichia Coli Periplasmic Histidine Binding Protein, HisJ. Prot. Sci. 26:268–279.

41. Feng, Y., L. Zhang, S. Wu, Z. Liu, X. Gao, X. Zhang, M. Liu, J. Liu, X. Huang, and W. Wang, 2016. Conformational Dynamics of Apo-GlnBP Revealed by Experimental and Computational Analysis. Angew. Chem. Int. Ed. 55:13990–13994.

42. Rambo, R. P., and J. A. Tainer, 2013. Accurate Assessment of Mass, Models and Resolution by Small-Angle Scattering. Nature 496:477–481.

43. Hura, G. L., H. Budworth, K. N. Dyer, R. P. Rambo, M. Hammel, C. T. McMurray, and J. A. Tainer, 2013. Comprehensive Objective Maps of Macromolecular Conformations by Quantitative SAXS Analysis. Nat Methods 10:453–454.

44. Semeraro, E. F., J. M. Devos, L. Porcar, V. T. Forsyth, and T. Narayanan, 2017. In Vivo Analysis of the Escherichia Coli Ultrastructure by Small-Angle Scattering. IUCrJ 4:751–757.

45. von Gundlach, A. R., V. M. Garamus, T. Gorniak, H. A. Davies, M. Reischl, R. Mikut, K. Hilpert, and A. Rosenhahn, 2016. Small Angle X-Ray Scattering as a High-Throughput Method to Classify Antimicrobial Modes of Action. Biochim. Biophys. Acta, Biomembr. 1858:918–925.

46. Watkin, S. A. J., T. M. Ryan, A. G. Miller, V. M. Nock, F. G. Pearce, and R. C. J. Dobson, 2017. Microfluidics for Small-Angle X-Ray Scattering. X-ray Scattering.

47. Pham, N., D. Radajewski, A. Round, M. Brennich, P. Pernot, B. Biscans, F. Bonneté, and S. Teychené, 2017. Coupling High Throughput Microfluidics and Small-Angle X-Ray Scattering to Study Protein Crystallization from Solution. Anal. Chem. 89:2282–2287.

48. Schwemmer, F., C. E. Blanchet, A. Spilotros, D. Kosse, S. Zehnle, H. D. T. Mertens, M. A. Graewert, M. Rössle, N. Paust, D. I. Svergun, F. von Stetten, R. Zengerle, and D. Mark, 2016. LabDisk for SAXS: A Centrifugal Microfluidic Sample Preparation Platform for Small-Angle X-Ray Scattering. Lab Chip 16:1161–1170.

