## Supplementary Materials for "Structure-based screening of binding affinities via small-angle X-ray scattering"

### Supplementary Information: Prediction of binding affinities from small-angle X-ray scattering titrations

[1](blank for initial submission)

#### Supplementary Methods

##### Computation of $V_\mathrm{R}$ in this study

The method proposed by Hura *et al.*, Nat. Methods, 2013 will be generally followed, with a small number of clarifications or modifications:

- Rather than setting a fixed molecular extent $D_{max}$ of 40 nm, an estimate of the protein molecular size is obtained by processing all HisBP SAXS curves with the DATGNOM utility. $D_{max}$ is thus set to 7.2 nm corresponding to the Shannon channel width of free HisBP.
- The binned Ratios $R$ are produced by conducting a geometrical-mean over individual values of $R(q)$.
- To account for additional scattering from unbound ligands and counterions that affect buffer subtraction, we further conduct Powell minimization over an added constant. Thus instead of computing $V_\mathrm{R}$ directly between two curves $I(q)$ versus $J(q)$, $V_\mathrm{R}$ is computed here for $I(q)-c$ versus $J(q)+c$ for a constant $c$ that minimizes $V_\mathrm{R}$. The symmetric subtraction/addition preserves the natural symmetry of $V_\mathrm{R}$.

We have implemented this procedure in python, available within the SAXScreen distribution under fit-saxs-curves.py.

$V_\mathrm{R}$ between two buffer-subtracted SAXS curves $I(q)$ and $J(q)$ over momentum transfer $q$ is computed as follows. First, the range of $q$ is pruned to exclude low-angle suffering from parasitic scattering and high-angle regions dominated by noise. The normalized ratio $R$ between the two curves are evaluated, ignoring any $q$ where either $I(q) < 0$ or $J(q) < 0$ so as to eliminate numerical instability:

$$R(q) = \frac{I(q)/J(q)}{\sum_q R(q)}, \tag{S1}$$

Shannon channel implies that $R(q)$ values within a local region of $q$-points are highly correlated and not independent, but can rather be split into N-degrees of freedom depending on molecular size. $R(q)$ is reduced according to the Shannon-channel width $\Delta q = \pi/D_{max}$, by applying a geometric mean over all $R(q)$ points within bins of width $\Delta q$. This produces an aggregate ratio $R_i$ for the $i$-th Shannon channel:

$$R_i = [R(q_j)R(q_{j+1})...R(q_{j+M-1})]^{1/M} \tag{S2}$$

M denotes the number of experimental $q$-points within each Shannon channel bin of width $\Delta q$, defined as a constant $N_{q,channel}$. In cases where $\Delta q$ does not evenly divide the input $q$-range, our implementation preserves a partial bin in the final sum of $V_\mathrm{R}$, but reduces its contribution to overall $V_\mathrm{R}$ by applying a fractional weight in the final sum. $V_{\mathrm{R}}$ is defined as the normalized differences between successive bins.

$$V_{\mathrm{R}} = 2w_i \sum_{i}^{N} \left\| \frac{R_i - R_{i+1}}{R_i + R_{i+1}} \right\|, \tag{S3}$$

$$w_i = \frac{M}{N_{q,channel}}. \tag{S4}$$

The weight $w_i$ evaluates to unity for all bins except for the last one at the highest angular region.

#### Supplementary Figures

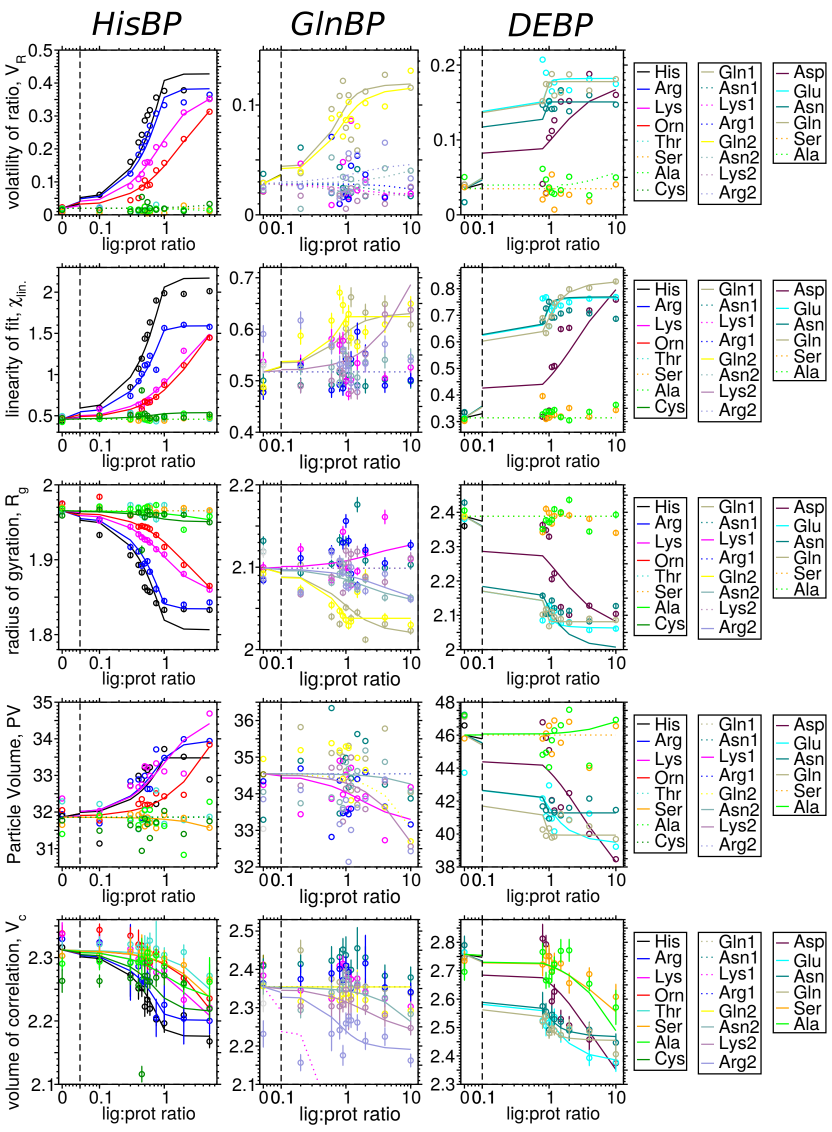


**Figure S1.** Titration of three periplasmic binding proteins at ESRF BM29 SAXS HisBP (left), GlnBP(center), and DEBP(right). Measured SAXS perturbations are expressed in a number of structural and scattering parameters, some of which form clear ligand-dependence. From top: volatility of ratio $V_{\mathrm{R}}$, linearity of fit $\chi_{\mathrm{lin.}}$, radius of gyration $R_{\mathrm{g}}$, particle volume, and volume of correlation $V_c$.

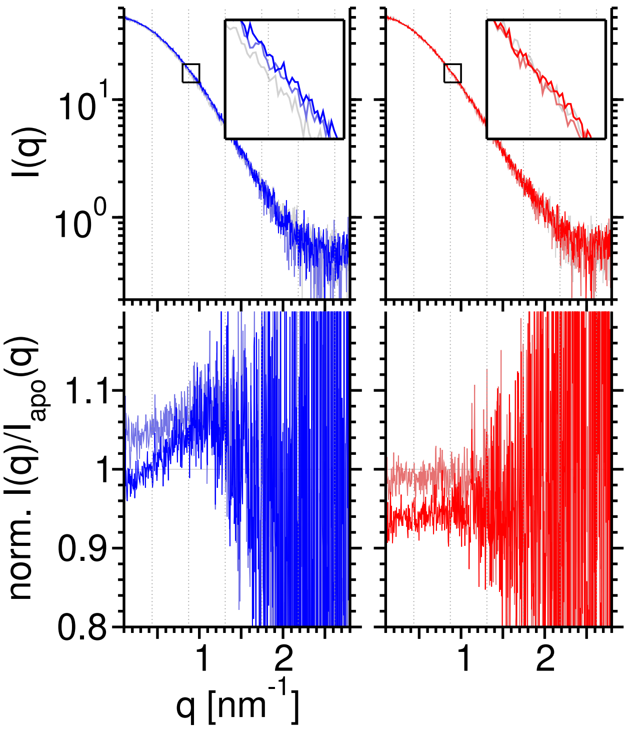


**Figure S2.** Scattering intensity of GlnBP:Gln (blue) and GlnBP:Asn(red), at three titration points apo (grey), 1:1 (light color) and 10:1 ligand-excess (color). Below each scattering curve are the normalized intensity ratios against respective apo-scattering.

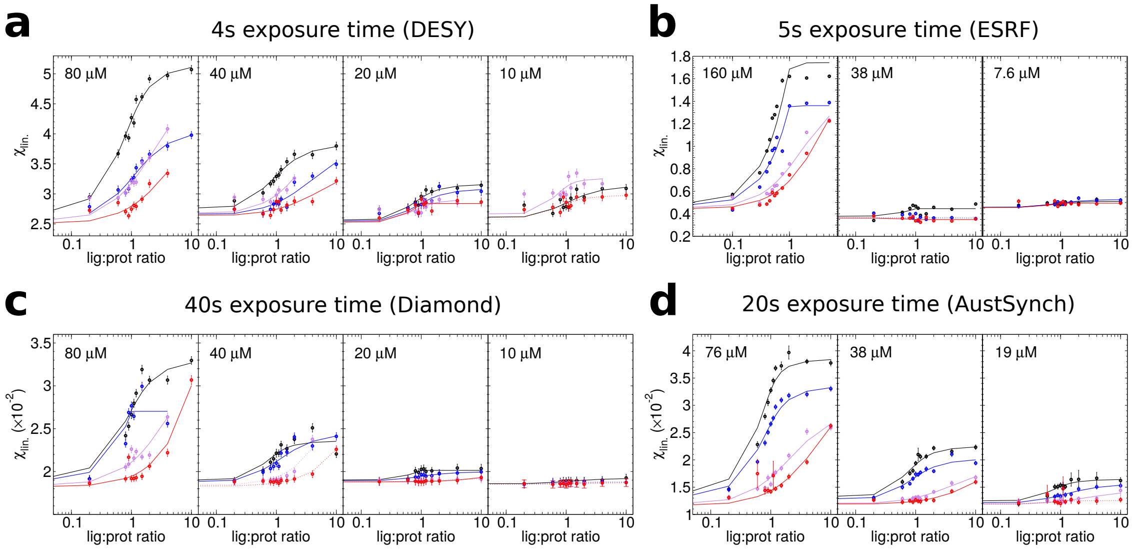


**Figure S3.** Replicate $\chi_{\rm lin.}$ titration curves of HisBP against four ligands His (black), Arg (blue), Lys (violet), and Orn (red), at DESY (a), ESRF (b), Diamond (c), and Australian Synchrotron (d). The HisBP concentrations used within each experiment are labelled on respective plots.

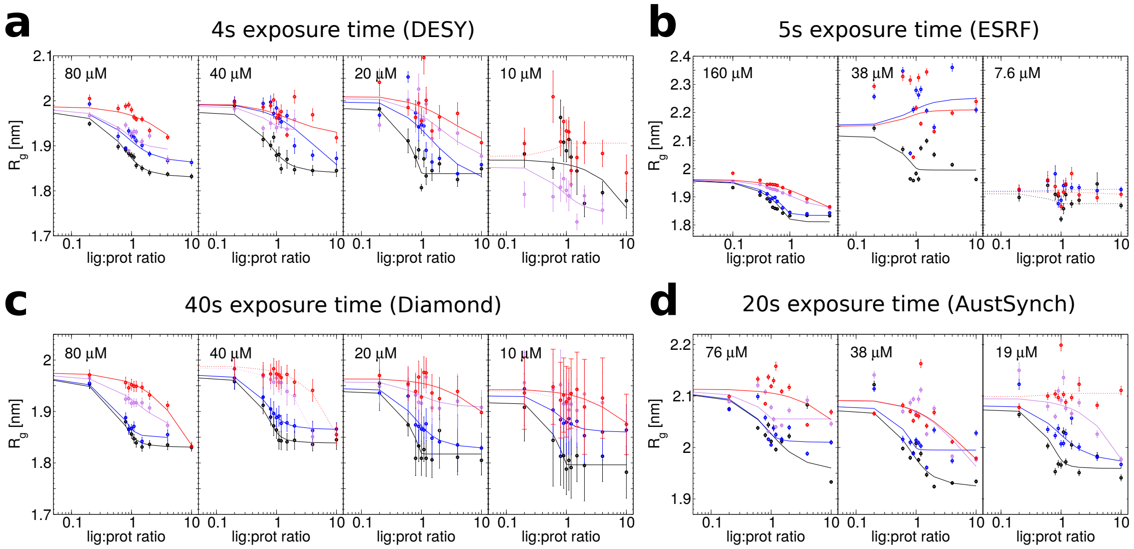


**Figure S4.** Replicate $R_\mathrm{g}$ titration curves of HisBP against four ligands His (black), Arg (blue), Lys (violet), and Orn (red), at DESY (a), ESRF (b), Diamond (c), and Australian Synchrotron (d). The HisBP concentrations used within each experiment are labelled on respective plots.

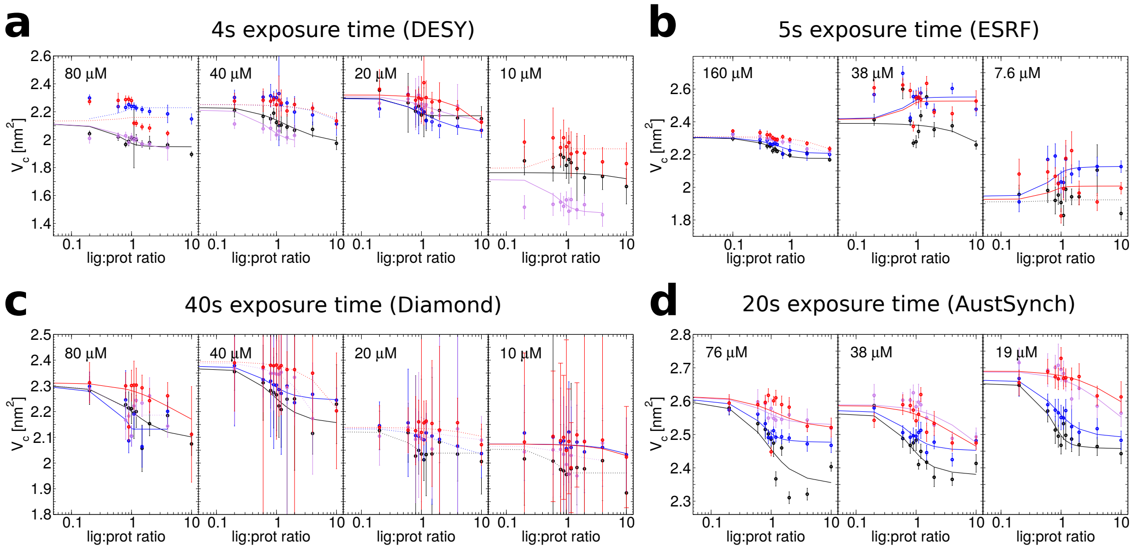


**Figure S5.** Replicate $V_c$ titration curves of HisBP against four ligands His (black), Arg (blue), Lys (violet), and Orn (red), at DESY (a), ESRF (b), Diamond (c), and Australian Synchrotron (d). The HisBP concentrations used within each experiment are labelled on respective plots.

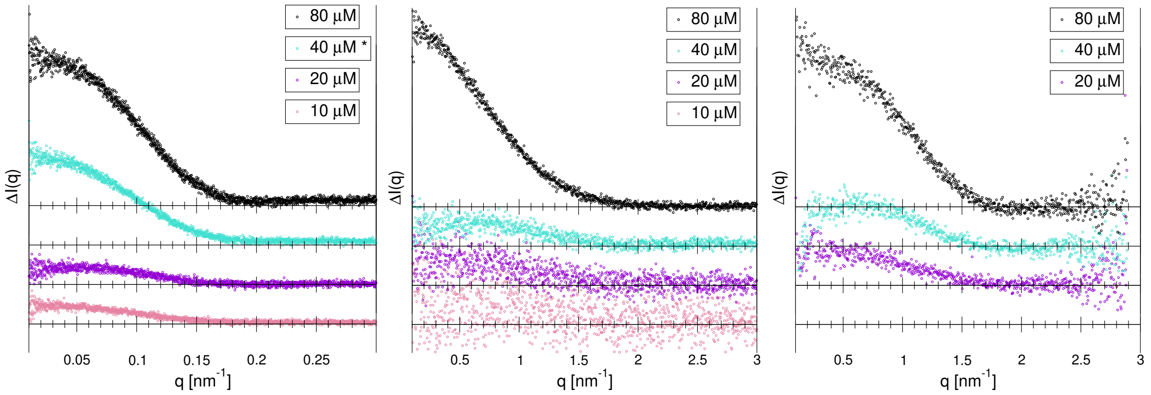


**Figure S6.** Difference spectra between His-saturated (10:1) and apo HisBP scattering curves as a function of protein concentration. Data collected at Diamond (left), DESY P12 (middle), and Australian Synchrotron SAXS/WAXS (right). The difference curve for His at Diamond, marked with a star, has been replaced with (4:1) measurement.

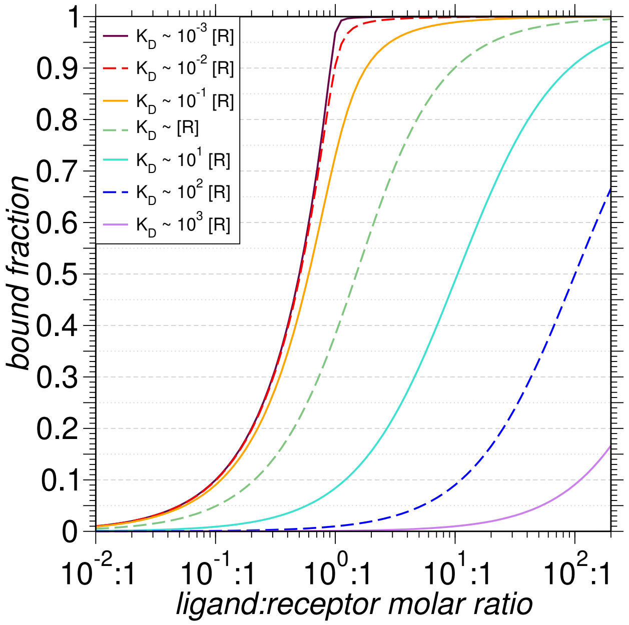


**Figure S7.** Theoretical two-state binding curve, scaled by the constant receptor concentration.

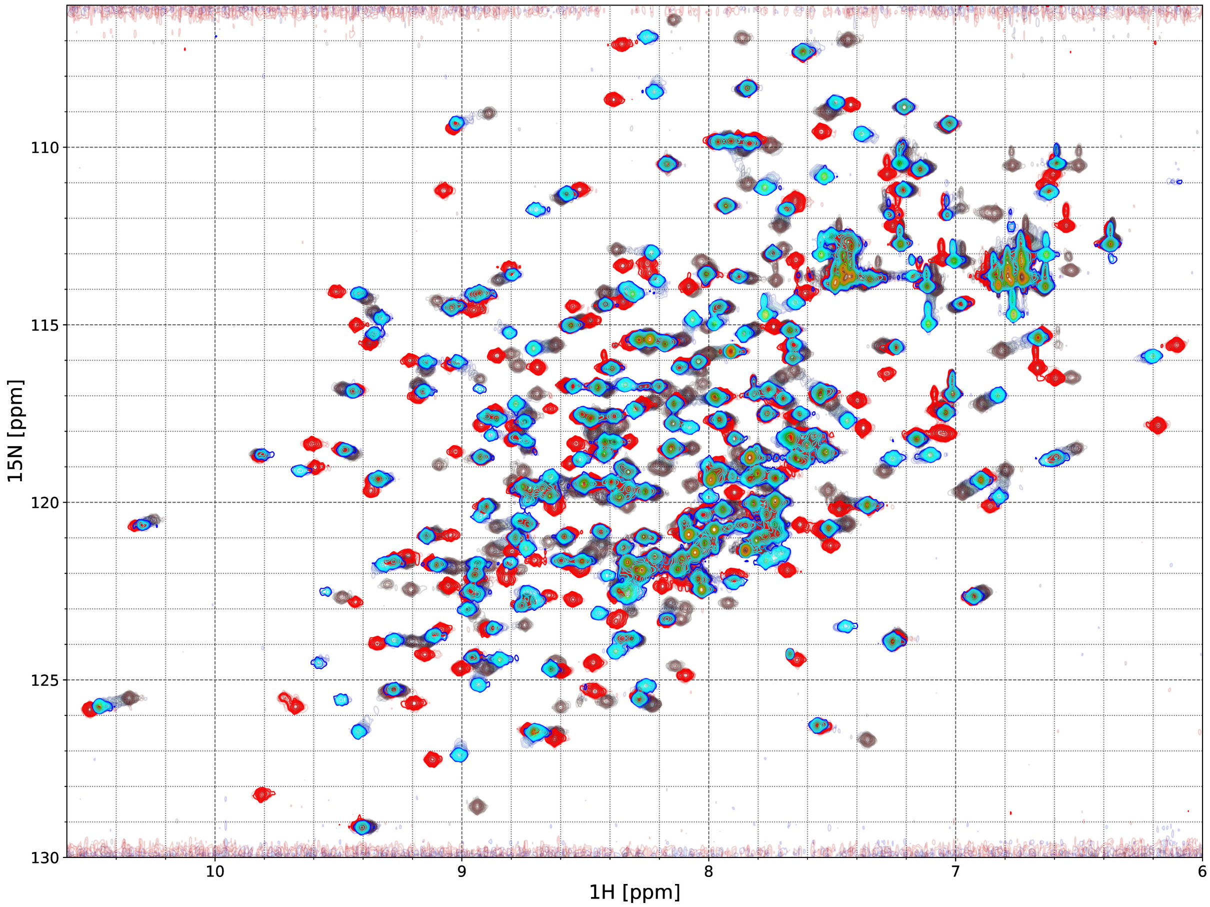


**Figure S8.** Titration of HisBP versus Histidine (red) and Arginine (blue-teal) by NMR, depicted by overlaying HSQC plots at multiple ligand:protein ratios between 0.0∼1.5. Resonance peaks of apo-HisBP are shown in grey, while resonance peaks from intermediate titration points are shown in light grey with gradually increasing color intensity until color saturation is reached at a ratio of 1.5:1.0. Partial differences in peak-shifts between HisBP:His and HisBP:Arg titrations indicate subtle differences in the final bound conformation.

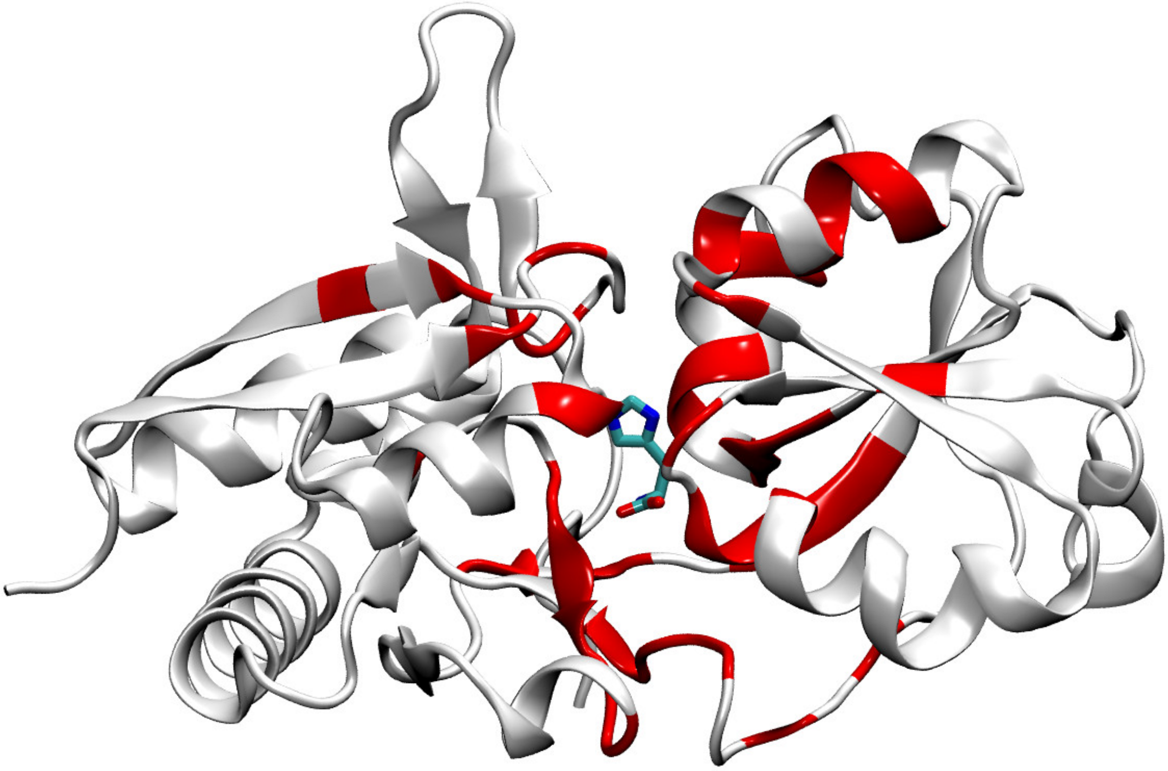

**Figure S9.** Depiction of HisBP crystal structure bound to Histidine (1HSL.pdb chain A), shown as white cartoons and sticks, respectively. Residues have been colored in red, where the $^{15}$N-$^{1}$H HSQC peaks of His-bound HisBP do not exhibit any overlap with peaks found in Arg-bound HisBP. Concentrations around the active-site cleft and inter-domain hinge suggest minor but extensive rearrangements to accommodate the larger Arg sidechain. Peak assignments taken from

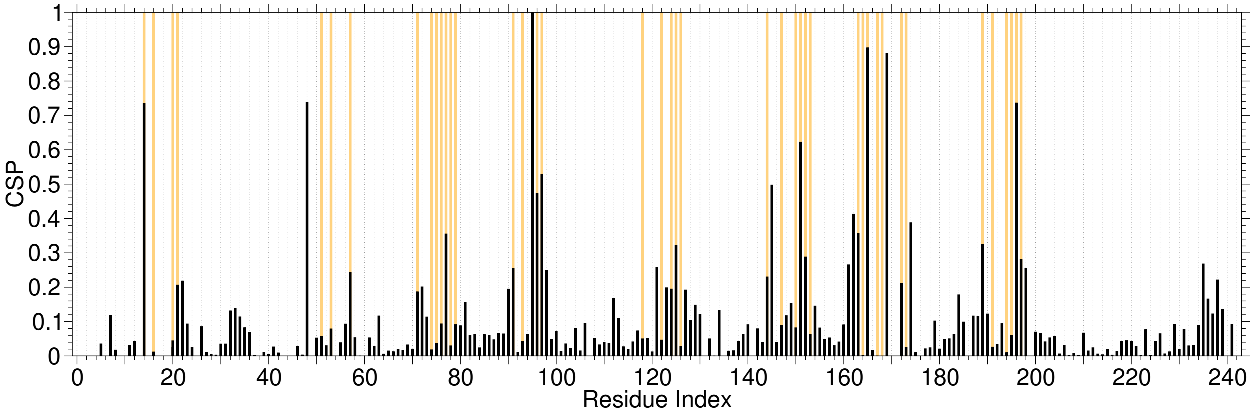


**Figure S10.** Residue-wise chemical-shift perturbations for Arg-bound HisBP relative to apo-HisBP (black). For comparison, residues highlighted in Figure S9 are shown here in orange.

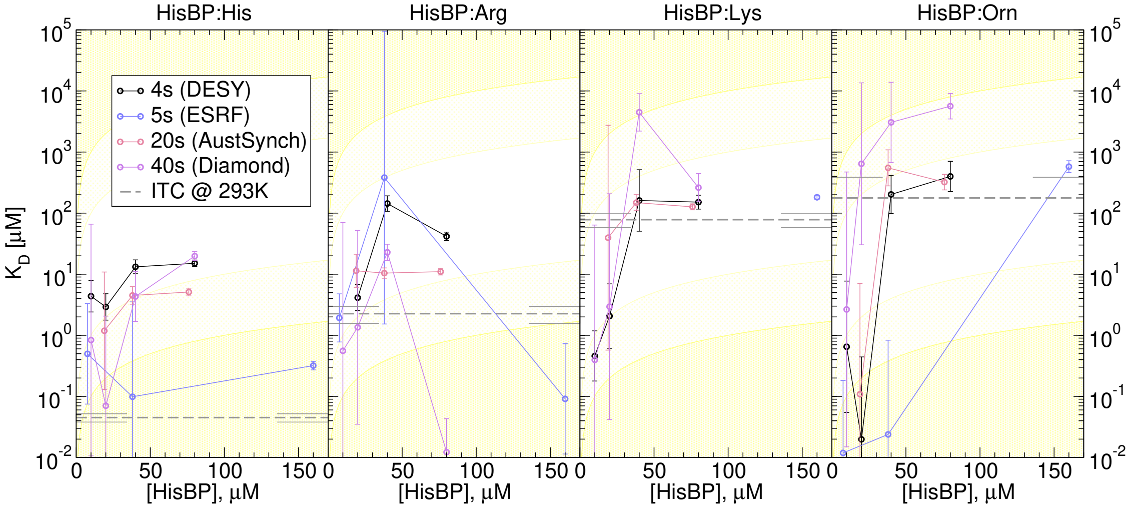


**Figure S11.** HisBP screens replicated at DESY P12 (black), Australian Synchrotron SAXS/WAXS (red), Diamond B21 (violet) and ESRF BM29 (blue), using $\chi_{\mathrm{lin.}}$ to derive relative populations. The affinity $K_D$ of HisBP versus four ligands is evaluated at different protein concentrations to test prediction capability versus ITC values (dotted grey with error bars). Regions where $K_D$ predictions lie outside 1 and 2 orders of magnitude of input protein concentrations are shaded in light and dark yellow, representing regions of decreased confidence.

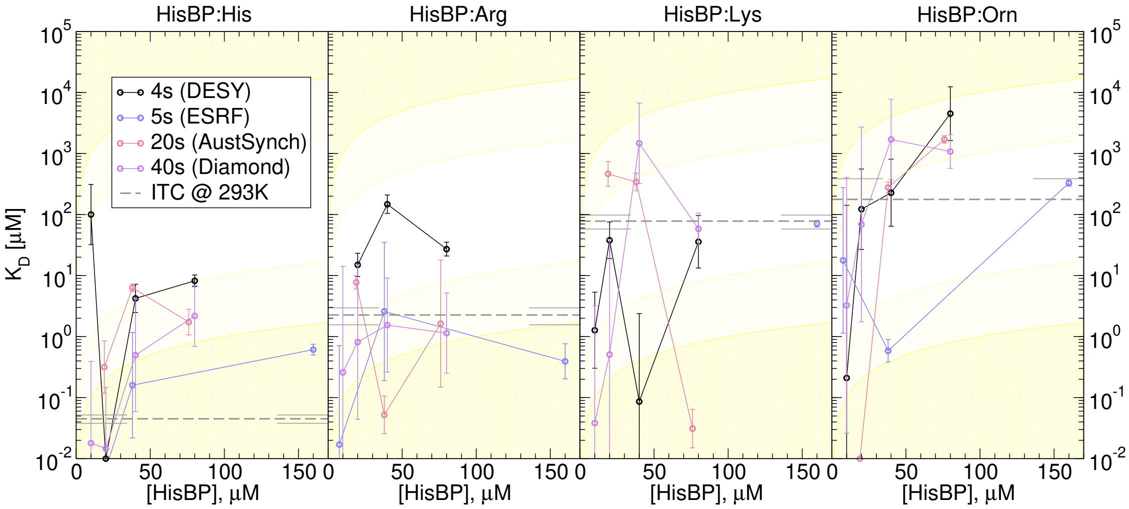


**Figure S12.** HisBP screens replicated at DESY P12 (black), Australian Synchrotron SAXS/WAXS (red), Diamond B21 (violet) and ESRF BM29 (blue), using $R_g$ to derive relative populations. The affinity $K_D$ of HisBP versus four ligands is evaluated at different protein concentrations to test prediction capability versus ITC values (dotted grey with error bars). Regions where $K_D$ predictions lie outside 1 and 2 orders of magnitude of input protein concentrations are shaded in light and dark yellow, representing regions of decreased confidence.

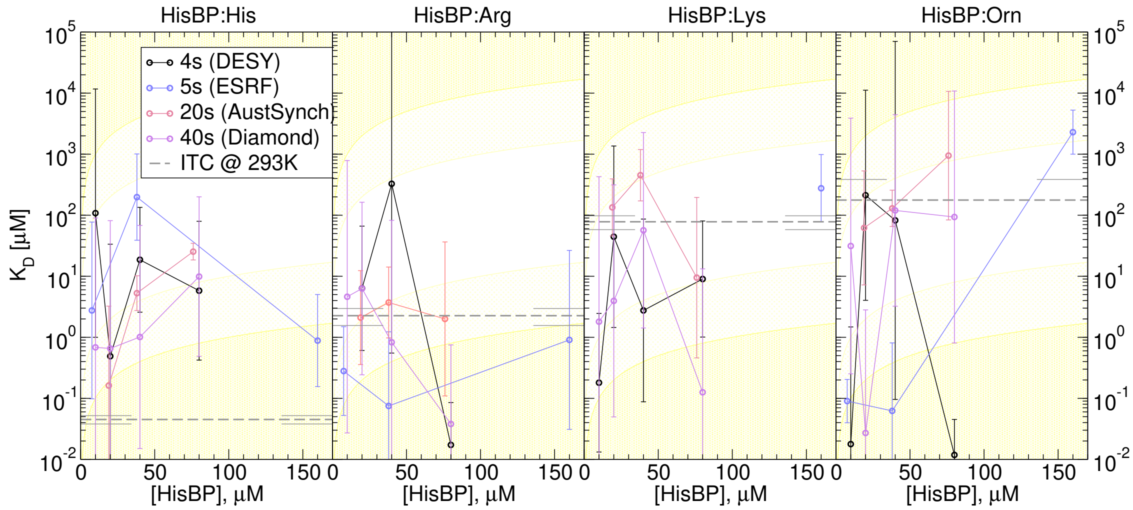


**Figure S13.** HisBP screens replicated at DESY P12 (black), Australian Synchrotron SAXS/WAXS (red), Diamond B21 (violet) and ESRF BM29 (blue), using $V_c$ to derive relative populations. The affinity $K_D$ of HisBP versus four ligands is evaluated at different protein concentrations to test prediction capability versus ITC values (dotted grey with error bars). Regions where $K_D$ predictions lie outside 1 and 2 orders of magnitude of input protein concentrations are shaded in light and dark yellow, representing regions of decreased confidence.

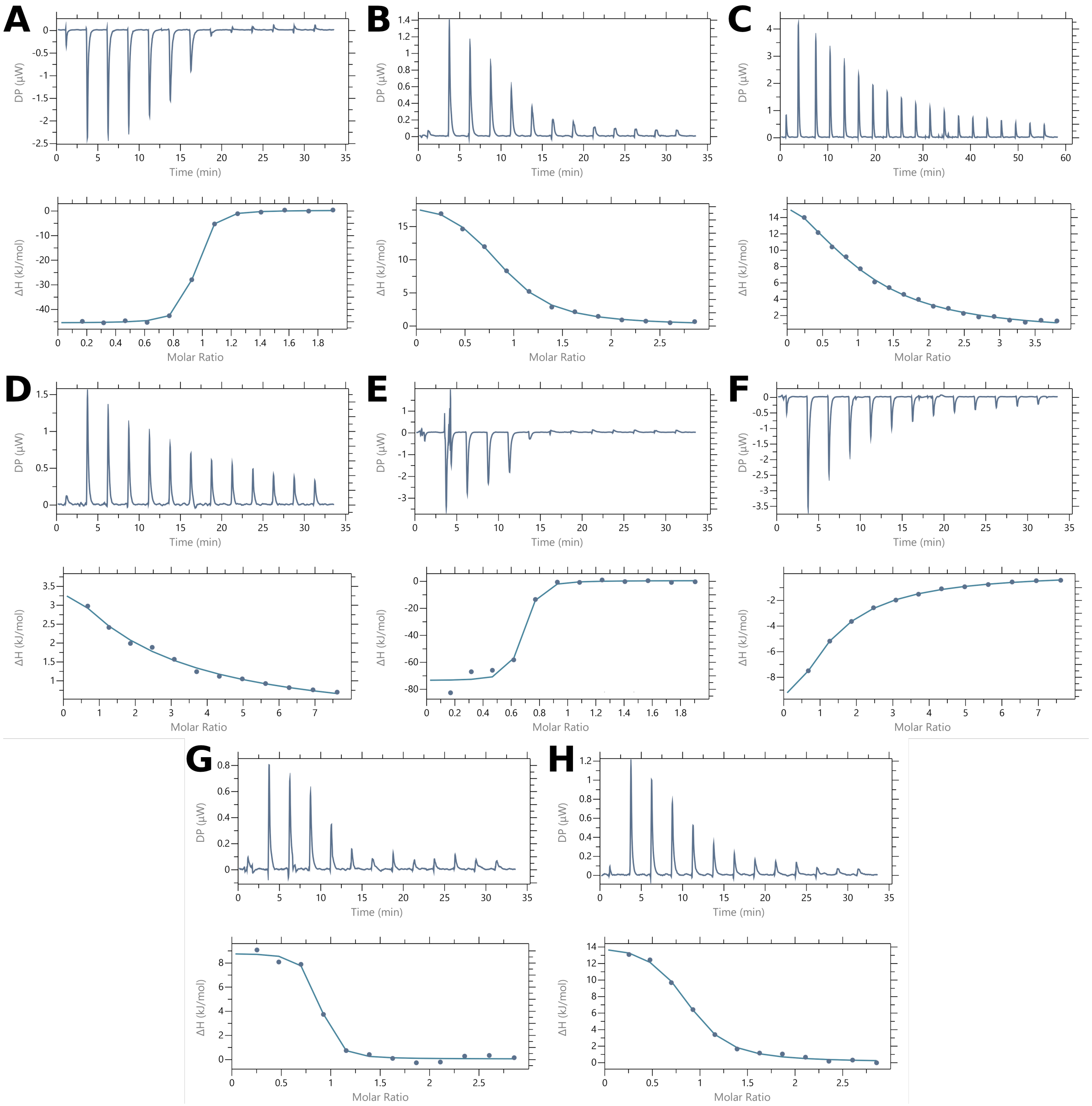


**Figure S14.** Titration curves of periplasmic binding proteins using isothermal calorimetry: (A) HisBP:His at 20 $\mu$M and 200 $\mu$M ; (B) HisBP:Arg at 20 $\mu$M and 300 $\mu$M ; (C) HisBP:Lys at 100 $\mu$M and 2 mM ; (D) HisBP:Orn at 50 $\mu$M and 2 mM ; (E) GlnBP:Gln at 20 $\mu$M and 200 $\mu$M ; (F) GlnBP:ARG at 50 $\mu$M and 2 mM ; (G) DEBP:Glu at 20 $\mu$M and 300 $\mu$M ; (H) DEBP:Asp at 20 $\mu$M and 300 $\mu$M.

#### Supplementary Tables

| Protein | Method | exposure | conc. ($\mu$M) | Ligand identity and dissociation constant $K_D$ | | | |
|---|---|---|---|---|---|---|---|
| | | | | His | Arg | Lys | Orn |
| HisBP | $V_{\mathrm{R}}$ (DESY) | 4 s | 80 | 6.4±1.1 $\mu$M | 5.9±0.6 $\mu$M | 58±7.5 $\mu$M | 100± 26 $\mu$M |
| | $V_{\mathrm{R}}$ (DESY) | 4 s | 40 | 3.7±0.58 $\mu$M | 74± 19 $\mu$M | 720±350 $\mu$M | 120± 43 $\mu$M |
| | $V_{\mathrm{R}}$ (DESY) | 4 s | 20 | 1±0.19 $\mu$M | 33±4.5 $\mu$M | 61±6.4 $\mu$M | 130±170 $\mu$M |
| | $V_{\mathrm{R}}$ (DESY) | 4 s | 10 | 84± 68 $\mu$M | n.d. | 2.3±1.6 nM | 130±2500 $\mu$M |
| | $V_{\mathrm{R}}$ (ESRF) | 5 s | 160 | 370±470 nM | 710±1000 nM | 49±4.1 $\mu$M | 250± 24 $\mu$M |
| | $V_{\mathrm{R}}$ (ESRF) | 5 s | 38 | 240±140 nM | 2±1.9 $\mu$M | n.d. | 5.4± 28 mM |
| | $V_{\mathrm{R}}$ (ESRF) | 5 s | 7.6 | 51±2900 $\mu$M | 2±190 $\mu$M | n.d. | 38± 53 $\mu$M |
| | $V_{\mathrm{R}}$ (AustSynch) | 20 s | 76 | 3.6±0.74 $\mu$M | 5.5±1.4 $\mu$M | 95± 11 $\mu$M | 220± 34 $\mu$M |
| | $V_{\mathrm{R}}$ (AustSynch) | 20 s | 38 | 1.5±0.7 $\mu$M | 3±1.4 $\mu$M | 32±4.6 $\mu$M | 190± 39 $\mu$M |
| | $V_{\mathrm{R}}$ (AustSynch) | 20 s | 19 | 62± 28 nM | 2.8±0.31 $\mu$M | 110± 10 $\mu$M | 130± 24 $\mu$M |
| | $V_{\mathrm{R}}$ (Diamond) | 40 s | 80 | 1.3±0.5 $\mu$M | 7.3±1.9 $\mu$M | 89±9.6 $\mu$M | 640±160 $\mu$M |
| | $V_{\mathrm{R}}$ (Diamond) | 40 s | 40 | 13± 11 nM | 4.8±0.48 $\mu$M | 320±210 $\mu$M | 680±360 $\mu$M |
| | $V_{\mathrm{R}}$ (Diamond) | 40 s | 20 | 5.1±4.1 nM | 920±110 nM | 47±3.6 $\mu$M | 1.4±1.1 mM |
| | $V_{\mathrm{R}}$ (Diamond) | 40 s | 10 | 1.4±0.92 nM | 370±140 nM | 230±200 $\mu$M | 1.2±1.8 mM |

**Table S1.** Raw fitted dissociation constant $K_D$ to SAXS $V_{\mathrm{R}}$ curves for all HisBP titrations. Noting that values outside of ∼2 orders of magnitudes of HisBP concentration are unreliable.

| Protein | Method | exposure | conc. ($\mu$M) | Ligand identity and dissociation constant $K_D$ | | | |
|---|---|---|---|---|---|---|---|
| | | | | His | Arg | Lys | Orn |
| HisBP | $\chi_{\mathrm{lin.}}$ (DESY) | 4 s | 80 | 15±1.9 $\mu$M | 42±6.7 $\mu$M | 150± 40 $\mu$M | 400±240 $\mu$M |
| | $\chi_{\mathrm{lin.}}$ (DESY) | 4 s | 40 | 13±3.5 $\mu$M | 140± 42 $\mu$M | 160±230 $\mu$M | 200±160 $\mu$M |
| | $\chi_{\mathrm{lin.}}$ (DESY) | 4 s | 20 | 2.9±1.5 $\mu$M | 4.1±2.1 $\mu$M | 2.1±3.2 $\mu$M | 20±220 nM |
| | $\chi_{\mathrm{lin.}}$ (DESY) | 4 s | 10 | 4.4±2.8 $\mu$M | n.d. | 460±500 nM | 650±3800 nM |
| | $\chi_{\mathrm{lin.}}$ (ESRF) | 5 s | 152 | 320± 51 nM | 91±360 nM | 180± 15 $\mu$M | 580±130 $\mu$M |
| | $\chi_{\mathrm{lin.}}$ (ESRF) | 5 s | 38 | 99±1800 nM | 380±47000 $\mu$M | n.d. | 24±420 nM |
| | $\chi_{\mathrm{lin.}}$ (ESRF) | 5 s | 7.6 | 500±1600 nM | 1.9± 2 $\mu$M | n.d. | 12± 91 nM |
| | $\chi_{\mathrm{lin.}}$ (AustSynch) | 20 s | 76 | 5.1±0.75 $\mu$M | 11±1.4 $\mu$M | 130± 14 $\mu$M | 320± 96 $\mu$M |
| | $\chi_{\mathrm{lin.}}$ (AustSynch) | 20 s | 38 | 4.5±1.5 $\mu$M | 10± 2 $\mu$M | 150± 46 $\mu$M | 550±400 $\mu$M |
| | $\chi_{\mathrm{lin.}}$ (AustSynch) | 20 s | 19 | 1.2±5.4 $\mu$M | 11±7.6 $\mu$M | 40±1400 $\mu$M | 110±3500 nM |
| | $\chi_{\mathrm{lin.}}$ (Diamond) | 40 s | 80 | 20±3.4 $\mu$M | 12± 20 nM | 260±150 $\mu$M | 5.6±2.8 mM |
| | $\chi_{\mathrm{lin.}}$ (Diamond) | 40 s | 40 | 4.3±4.7 $\mu$M | 23±7.1 $\mu$M | 4.5±3.4 mM | 3.1±6.6 mM |
| | $\chi_{\mathrm{lin.}}$ (Diamond) | 40 s | 20 | 70±1000 nM | 1.4± 26 $\mu$M | 2.9±100 $\mu$M | 640±6800 $\mu$M |
| | $\chi_{\mathrm{lin.}}$ (Diamond) | 40 s | 10 | 840±33000 nM | 560±35000 nM | 400±32000 nM | 2.7±240 $\mu$M |

**Table S2.** Raw fitted dissociation constant $K_D$ to SAXS $\chi_{\mathrm{lin.}}$ curves for all HisBP titrations. Noting that values outside of ∼2 orders of magnitudes of HisBP concentration are unreliable.

| Protein | Method | exposure | conc. ($\mu$M) | Ligand identity and dissociation constant $K_D$ | | | |
|---|---|---|---|---|---|---|---|
| | | | | His | Arg | Lys | Orn |
| HisBP | $R_g$ (DESY) | 4 s | 80 | 8.2±1.8 $\mu$M | 27±7.1 $\mu$M | 36± 42 $\mu$M | 4.5±5.4 mM |
| | $R_g$ (DESY) | 4 s | 40 | 4.2±2.4 $\mu$M | 150± 52 $\mu$M | 86±1200 nM | 230±370 $\mu$M |
| | $R_g$ (DESY) | 4 s | 20 | 2.7±4.7 nM | 15±6.8 $\mu$M | 38± 28 $\mu$M | 120±270 $\mu$M |
| | $R_g$ (DESY) | 4 s | 10 | 100±140 $\mu$M | n.d. | 1.3±2.5 $\mu$M | 210±71000 nM |
| | $R_g$ (ESRF) | 5 s | 160 | 610±120 nM | 390±280 nM | 71±8.9 $\mu$M | 330± 38 $\mu$M |
| | $R_g$ (ESRF) | 5 s | 38 | 160±570 nM | 2.6± 17 $\mu$M | n.d. | 580±250 nM |
| | $R_g$ (ESRF) | 5 s | 7.6 | 0.76±2.1e-05 nM | 17±360 nM | n.d. | 18±140 $\mu$M |
| | $R_g$ (AustSynch) | 20 s | 76 | 1.7±0.87 $\mu$M | 1.6±8.8 $\mu$M | 31± 25 nM | 1.7±0.24 mM |
| | $R_g$ (AustSynch) | 20 s | 38 | 6.3±0.87 $\mu$M | 52± 40 nM | 340±120 $\mu$M | 280± 58 $\mu$M |
| | $R_g$ (AustSynch) | 20 s | 19 | 320±370 nM | 7.7± 2 $\mu$M | 460±230 $\mu$M | 2.3±3.9 nM |
| | $R_g$ (Diamond) | 40 s | 80 | 2.2±3.1 $\mu$M | 1.1±2.5 $\mu$M | 58± 37 $\mu$M | 1.1±0.73 mM |
| | $R_g$ (Diamond) | 40 s | 40 | 500±2100 nM | 1.5±4.4 $\mu$M | 1.5±3.2 mM | 1.7±3.7 mM |
| | $R_g$ (Diamond) | 40 s | 20 | 15± 73 nM | 810±7500 nM | 510±20000 nM | 69±1400 $\mu$M |
| | $R_g$ (Diamond) | 40 s | 10 | 18±200 nM | 260±7000 nM | 38±1600 nM | 3.3±200 $\mu$M |

**Table S3.** Raw fitted dissociation constant $K_D$ to SAXS $R_g$ curves for all HisBP titrations. Noting that values outside of ~2 orders of magnitudes of HisBP concentration are unreliable.

| Protein | Method | exposure | conc. ($\mu$M) | Ligand identity and dissociation constant $K_D$ | | | |
|---|---|---|---|---|---|---|---|
| | | | | His | Arg | Lys | Orn |
| HisBP | $V_c$ (DESY) | 4 s | 80 | 5.8± 40 $\mu$M | 17± 41 nM | 9± 40 $\mu$M | 12± 21 nM |
| | $V_c$ (DESY) | 4 s | 40 | 19± 66 $\mu$M | 330±97000 $\mu$M | 2.8± 43 $\mu$M | 82±35000 $\mu$M |
| | $V_c$ (DESY) | 4 s | 20 | 490±17000 nM | 6.3± 33 $\mu$M | 44±680 $\mu$M | 210±5600 $\mu$M |
| | $V_c$ (DESY) | 4 s | 10 | 110±5900 $\mu$M | n.d. | 180±1200 nM | 18±740 nM |
| | $V_c$ (ESRF) | 5 s | 160 | 880±2400 nM | 910±13000 nM | 280±450 $\mu$M | 2.3±2.1 mM |
| | $V_c$ (ESRF) | 5 s | 38 | 200±490 $\mu$M | 75±610 nM | n.d. | 62±400 nM |
| | $V_c$ (ESRF) | 5 s | 7.6 | 2.8± 39 $\mu$M | 280±720 nM | n.d. | 90± 82 nM |
| | $V_c$ (AustSynch) | 20 s | 76 | 25±8.1 $\mu$M | 2± 18 $\mu$M | 9.5± 98 $\mu$M | 950±5300 $\mu$M |
| | $V_c$ (AustSynch) | 20 s | 38 | 5.3±3.7 $\mu$M | 3.7±6.6 $\mu$M | 450±510 $\mu$M | 130± 96 $\mu$M |
| | $V_c$ (AustSynch) | 20 s | 19 | 160±1600 nM | 2.1± 6 $\mu$M | 130±170 $\mu$M | 62±260 $\mu$M |
| | $V_c$ (Diamond) | 40 s | 80 | 9.9±100 $\mu$M | 38±380 nM | 130±6600 nM | 94±5400 $\mu$M |
| | $V_c$ (Diamond) | 40 s | 40 | 1± 34 $\mu$M | 830±41000 nM | 57±1100 $\mu$M | 120±2200 $\mu$M |
| | $V_c$ (Diamond) | 40 s | 20 | 660±41000 nM | 6.3± 81 $\mu$M | 3.9±160 $\mu$M | 27±1400 nM |
| | $V_c$ (Diamond) | 40 s | 10 | 680±45000 nM | 4.6±390 $\mu$M | 1.8±210 $\mu$M | 31±1900 $\mu$M |

**Table S4.** Raw fitted dissociation constant $K_D$ to SAXS $V_c$ curves for all HisBP titrations. Noting that values outside of ~2 orders of magnitudes of HisBP concentration are unreliable.

**Table S5.** Detailed reporting for SAXS measurements for six samples, measured at ESRF BM29 during Sep. 2018: (i) HisBP in free state and bound to ten-fold molar excess His, (ii) GlnBP in free state and bound to ten-fold molar excess Gln, amd (iii) DEBP in free state and bound to ten-fold molar excess Glu.

| | $HisBP_{free}$ | $HisBP_{bound}$ | $GlnBP_{free}$ | $GlnBP_{bound}$ | $DEBP_{free}$ | $DEBP_{bound}$ |
|---|---|---|---|---|---|---|
| **(a) Sample Details** | | | | | | |
| Source organism | *E. coli* | | | | | |
| Expression organism | *E. coli* BL21(DE3) | | | | | |
| Plasmid source | this work | | Prof. Colin Jackson | | | |
| Description | P0AEU0 (23-260) with N-terminal Gly | | P0AEQ3 (23-248) with N-terminal $His_6$ | | P37902 (28-302) with N-terminal $His_6$ | |
| Computed extinction coefficient $\epsilon_{280nm}$ ($M^{-1}cm^{-1}$) | 17,545 | | 25,900 | | 24,075 | |
| Molecular mass $M$ from chemical composition (kDa) | 26.290 | | 25.786 | | 32.052 | |
| loading concentration ($mg\,ml^{-1}$) | 4.2 | | 2.0 | | 2.2 | |
| injection volume ($\mu$l) | 50 | | 50 | | 50 | |
| Concentration ($\mu$M) | 160 | | 80 | | 70 | |
| Solvent composition and source | 100 mM NaCl, 20 mM $NaPO_4$, 0.5 mM TCEP, pH 7.4 | | | | | |
| **(b) SAS data collection parameters** | | | | | | |
| Source and instrument | Grenoble ESRF BM29 with Dectris Pilatus 1M | | | | | |
| Wavelength (Å) | 0.9919 | | | | | |
| Sample-detector distance (m) | 2.849 | | | | | |
| $q$-measurement range ($nm^{-1}$) | 0.0306–4.9462 | | | | | |
| Intensity Normalization | 0.00192 | | | | | |
| Radiation damage monitoring | frame-by-frame comparison | | | | | |
| Exposure time (s) & number | $2.0 \times 12$ frames | | | | | |
| Sample configuration | 96-well plate with flow-through capillary measurement | | | | | |
| Sample temperature (°C) | 20 | | | | | |
| **(c) Software employed for SAS data reduction, analysis and interpretation** | | | | | | |
| SAXS data processing | SAXScreen and ATSAS 2.8 | | | | | |
| Molecular graphics | Visual Molecular Dynamics | | | | | |
| **(d) Structural parameters** | | | | | | |
| Guinier analysis (PRIMUS) | | | | | | |
| $I(0)$ (raw) | 124.42±0.10 | 128.71±0.10 | 49.40±0.08 | 49.65±0.06 | 67.04±0.11 | 67.06±0.08 |
| $R_g$ (nm) | 1.96±0.07 | 1.83 ±0.10 | 2.11±0.14 | 2.03±0.13 | 2.32±0.21 | 2.06±0.13 |
| $q$-range ($nm^{-1}$) | 0.1249–0.6580 | 0.0825–0.7052 | 0.0919–0.6156 | 0.0966–0.6391 | 0.0872–0.5495 | 0.1297–0.6297 |
| Coefficient of correl. $R^2$ | 0.97 | 0.99 | 0.94 | 0.95 | 0.96 | 0.95 |
| $M$ from $I(0)$ (kDa, ratio to expected value) | 29.6 (1.13) | n.a. | 24.7 (0.96) | n.a. | 30.47 (0.95) | n.a. |
| $P(r)$ Analysis (AUTOGNOM) | | | | | | |
| $I(0)$ ($cm^{-1}$) | 124.7±0.1 | 128.8±0.1 | 49.13±0.09 | 49.47±0.09 | 67.28±0.12 | 66.97±0.10 |
| $R_g$ (nm) | 1.975±0.002 | 1.833±0.002 | 2.097±0.005 | 2.023±0.004 | 2.380±0.008 | 2.061±0.005 |
| $d_{max}$ (nm) | 6.03 | 5.71 | 6.18 | 6.10 | 8.46 | 6.44 |
| $q$-range ($nm^{-1}$) | 0.1014–2.7998 | | | | | |
| GNOM total est. | 0.9759 | 0.9691 | 0.8962 | 0.8913 | 0.7415 | 0.8793 |
| $M$ from $I(0)$ (ratio to expected value) | 29.7 (1.13) | n.a. | 24.6 (0.95) | n.a. | 30.6 (0.95) | n.a. |

| | $HisBP_{free}$ | $HisBP_{bound}$ | $GlnBP_{free}$ | $GlnBP_{bound}$ | $DEBP_{free}$ | $DEBP_{bound}$ |
|---|---|---|---|---|---|---|
| (g) Data and model deposition IDs | | | | | | |
| SASBDB | SASDFD8 | SASDFE8 | SASDFF8 | SASDFG8 | SASDFH8 | SASDFJ8 |
